# Hippo signalling regulates the nuclear behaviour and DNA dwell times of YAP and TEAD to control transcription

**DOI:** 10.1101/2025.03.11.642705

**Authors:** Benjamin Kroeger, Samuel A. Manning, Varshini Mohan, Qiji Deng, Jieqiong Lou, Guizhi Sun, Sara Lamont, Alex J. McCann, Mathias Francois, Jose M. Polo, Elizabeth Hinde, Kieran F. Harvey

## Abstract

Over the past two decades, genetic and proteomic screens have enabled the discovery and elucidation of the Hippo pathway as a complex signalling network that controls tissue growth and cell fate and which is of major importance for human cancers. Despite these advances, our understanding of how Hippo signalling regulates transcription is less clear. To address this, we used live microscopy approaches to study the nuclear behaviour of the major transcription effectors of the human Hippo pathway, YAP and TEADs. Our experiments revealed that TEADs are a major determinant of YAP’s nuclear biophysical behaviour, whilst YAP only has a minor influence on TEAD behaviour. Acute chemical inhibition of Hippo signalling stimulated an increase in the DNA residence time of both YAP and TEAD1. Consistently, YAP and TEAD1 bound DNA for longer periods in cells with high intrinsic YAP/TEAD activity (induced trophoblast stem cells) than in cells with low intrinsic YAP/TEAD activity (induced pluripotent stem cells). TEAD1 bound the genome on a broad range of timescales, and this is extended substantially in nuclear condensates. Finally, single molecule tracking experiments revealed that a fusion protein encoded by a cancer-associated YAP allele exhibits substantially different nuclear biophysical behaviour than either YAP or TEAD1. These live microscopy experiments reveal that Hippo signalling regulates transcription at least in part by influencing the DNA dwell times of both YAP and TEAD.

## INTRODUCTION

The Hippo pathway is an evolutionarily conserved signalling network that has been implicated in the control of organ growth and regeneration, as well as specific cell fate decisions (*1–4*). It operates in a wide range of species, from single celled metazoan ancestors to humans, and from the earliest stages of mammalian development through to homeostasis in adults. Defective Hippo signalling also underpins different human diseases, including several cancers (*5*). The Hippo pathway was first discovered and elucidated using *Drosophila melanogaster* genetic screens. For simplicity, here, we will use the human nomenclature for Hippo pathway proteins, given our study is based in human cells. The central DNA binding transcription factors in the Hippo pathway are TEAD1-TEAD4. In partnership with the YAP and TAZ transcription co-activators, TEADs can promote transcription of their target genes (*6–9*). Additionally, TEADs can repress transcription with either VGLL4 or INSM1A, both of which can also form a physical complex with TEADs (*10–13*). YAP and TAZ activity are regulated by the Hippo pathway core kinase cassette, which in turn are regulated by a range of upstream signalling proteins (*1–4*). These proteins respond to a variety of cues, including cell-cell adhesion, cell polarity, mechanical forces and cellular stresses associated with changes in energy availability and osmolarity (*1–4*).

To date, the best defined regulatory process in YAP/TAZ-mediated transcription is control of the nucleo-cytoplasmic shuttling rate of YAP and TAZ by the Hippo pathway (*14*). The central Hippo pathway kinases LATS1/2 phosphorylate YAP/TAZ on multiple serine residues, which limits the nuclear pool of these proteins and their ability to induce transcription with the TEADs (*15–18*). LATS-mediated phosphorylation was initially thought to cause stable sequestration of YAP/TAZ in the cytoplasm, although live imaging studies revealed that the majority of YAP and TAZ continually and rapidly shuttle between the cytoplasm and nucleus, and both Hippo signalling and TEAD abundance modulate the rate at which this happens (*19–23*). In addition, Hippo pathway-independent processes can influence YAP/TAZ nucleocytoplasmic shuttling such as nuclear deformation, mechanical forces and phosphorylation by additional kinases (*20, 24, 25*). Furthermore, YAP and TAZ can both undergo liquid-liquid phase separation, and their transcription-promoting activity is higher in YAP/TAZ-enriched nuclear condensates than other regions of the nucleus (*26–29*).

What is currently unclear is whether the sole mechanism by which the Hippo pathway regulates transcription is to dictate the YAP/TAZ nucleocytoplasmic shuttling rate, or whether it does so by additional mechanisms. Recently, live microscopy experiments in *Drosophila* tissues revealed that Yki (YAP/TAZ orthologue) overexpression extends Scalloped (TEAD) DNA binding times, whilst overexpression of the transcription co-repressors Tgi (VGLL4) and Nerfin-1 (INSM1) had the opposite effect. However, it is unknown whether this is also true in human cells, or whether changes in Hippo signalling alter YAP/TEAD biophysical behaviour and/or DNA dwell times. Further, alterations in transcription factor DNA dwell times have been associated with both transcription activation and repression. For example, increased DNA dwell times of the Serum Response Factor transcription factor are observed when transcription is elevated (*30*), whilst extended DNA binding of transcription repressors has been show to correlate with reduced transcription (*31*).

To investigate the outstanding question of how Hippo signalling regulates transcription, we employed high-resolution live imaging modalities to assess the biophysical behaviour of YAP and TEAD1 in cells with either intrinsic or chemically induced differences in Hippo pathway activity, as well as in nuclear condensates. Finally, to gain insights into the importance of biophysical behaviour for transcription factor function, we investigated a cancer-associated variant of YAP (YAP-TFE3) which drives the sarcoma epithelioid hemangioendothelioma (*32, 33*).

## RESULTS

### YAP and TEAD1 exhibit different biophysical properties in cell nuclei

To investigate the biophysical behaviour of the transcription effectors of the Hippo pathway, YAP and TEADs, we employed two microscopy techniques: single molecule tracking (SMT) and fluorescence fluctuation spectroscopy. To assess YAP and TEAD behaviour at single molecule resolution we generated plasmids that expressed versions of YAP or TEAD1 fused to HaloTag. HaloTag is derived from a bacterial enzyme that covalently binds to a fluorescently tagged ligand that is bright, highly resistant to photobleaching, and is amenable to SMT (*34*). We imaged YAP and TEAD1 at single molecule resolution and tracked their behaviour in the nucleus of living cells, using highly inclined and laminated optical sheet (HILO) microscopy, which is a variant of total internal reflection fluorescence (TIRF) microscopy (*35*). Most experiments were performed using doxycycline-inducible transgenes that were stably integrated into the genome of human MCF10A breast epithelial cells, which are immortalised but non-transformed. Doxycycline was titrated to induce expression of HaloTagged YAP and TEAD1 at endogenous levels or lower (fig. S1, a and b).

Initially, we acquired images of cells every 20ms (fast SMT) to capture the behaviour of the majority of TEAD1 and YAP molecules (Fig. 1, A and B, and movie S1). Using PALMTracer we tracked the trajectories of each TEAD1 and YAP molecule and plotted their molecular mobilities (*36*) Both proteins displayed a bimodal mobility distribution comprised of mobile and immobile molecules, with YAP in general being more mobile than TEAD1 (Fig. 1C). This was confirmed by calculating the mean squared displacement of individual TEAD1 and YAP proteins (Fig. 1D), the ratio of mobile to immobile molecules (Fig. 1E) and the area under the curve (the distance a population of molecules moves in a given time) (Fig. 1F). These different nuclear mobilities likely represent proteins that are freely diffusing (mobile) or bound to DNA (immobile) (*34*). HaloTagged TEAD1 behaviour did not vary significantly when expressed at different levels, whilst HaloTagged YAP became more mobile when expressed approximately ten-fold higher than endogenous YAP (fig. S1c to g), highlighting the importance of studying proteins expressed at their native levels, which was done for most subsequent experiments. We also found that the position of the HaloTag (N-terminal or C-terminal) did not have a major influence on nuclear TEAD1 or YAP behaviour (fig. S2, a to d).

**Figure 1.**
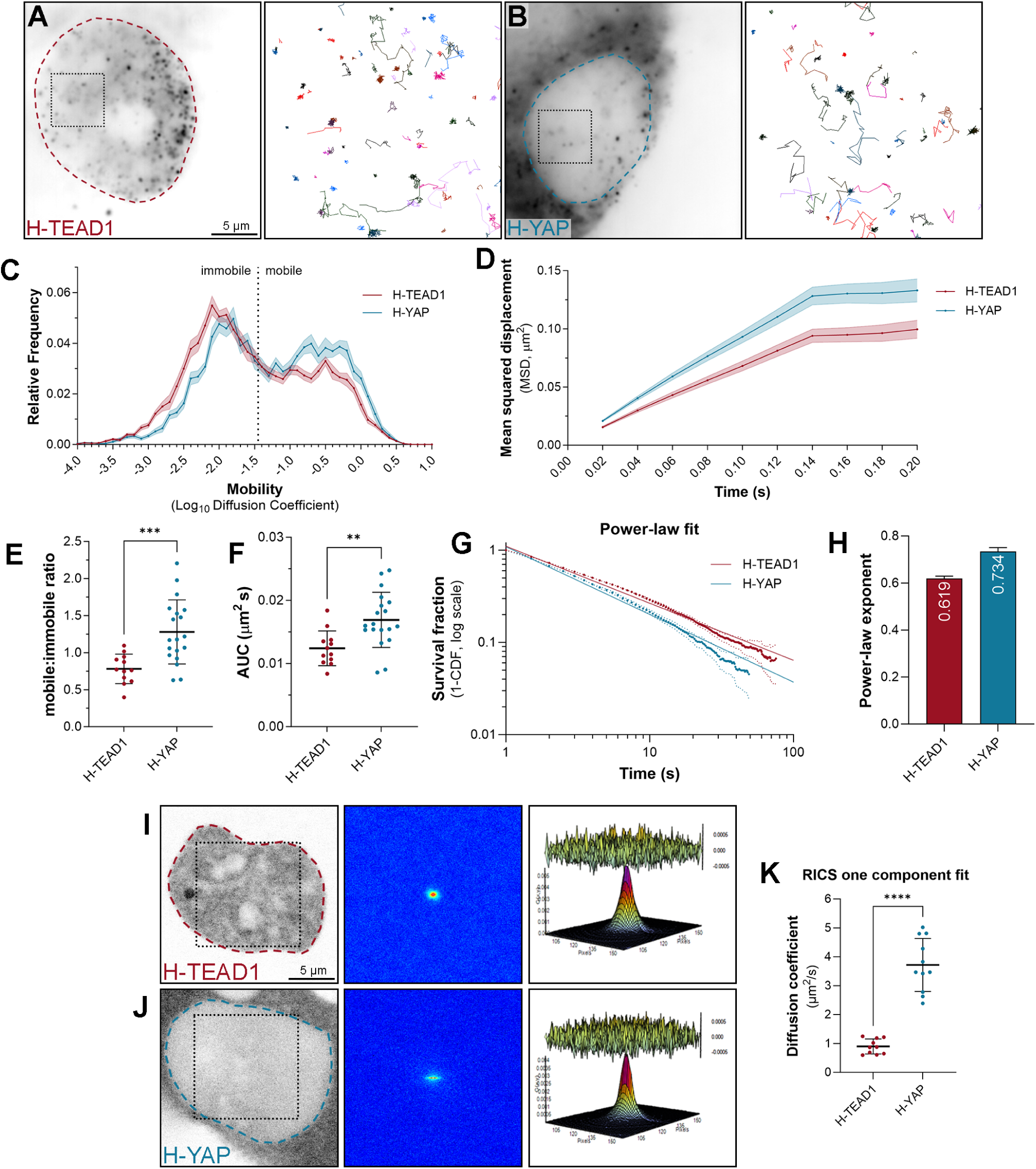
YAP and TEAD1 exhibit different biophysical properties in cell nuclei. **A and B)** Left panels are average intensity projection images of single MCF10A cells expressing Halo-tagged TEAD1 (A) or YAP (B), imaged at single molecule resolution by HILO. Dashed outlines indicate the nucleus. Right panels show the trajectories of individual TEAD1, or YAP molecules tracked over time. Boxed regions in cell images indicate the area corresponding to trajectories. Scale bar is indicated. **C**) Chart showing the relative frequency of molecule mobilities (Log_10_ diffusion coefficient) for TEAD1 (red) and YAP (blue) molecules in MCF10A cell nuclei over time. Data presented as the mean ± SEM. Mobile and immobile fractions are indicated. Data from 14,053 trajectories from 12 cells for TEAD1 and 6,386 trajectories from 19 cells for YAP. **D**) Chart showing the mean squared displacement (µm^2^) of TEAD1 (red) and YAP (blue) molecules in MCF10A cell nuclei over time. Data presented as the mean ± SEM. **E and F)** Charts showing the mobile to immobile ratio (E) or the area under the curve (F, µm^2^/s) of TEAD1 and YAP molecules in MCF10A cell nuclei. Data presented as mean ± SD, p-values obtained using an unpaired t-test; *** p < 0.001, ** p < 0.01, n = 12 for TEAD1 and 19 for YAP. **G)** Chart showing the photobleach-corrected survival distribution of single Halo-TEAD1 or Halo-YAP molecules in the nucleus of MCF10A cells, with power-law fits (solid lines). Dotted lines are ± 99% CI. n = 61,860 trajectories from 31 cells for TEAD1, and 27,748 trajectories from 20 cells for YAP. **H)** Bar chart showing the power-law exponents of TEAD1 and YAP in MCF10A cells. Error bars indicate 95% CI. **I-J)** Left panels are confocal microscope images of TEAD1 (H) or YAP (I) in single MCF10A cells. Scale bar is indicated. Boxed region shows analysed nuclear region. Central panels are three-dimensional RICS correlation functions calculated from the image acquisitions. Right panel shows the single-component diffusion model fit of the 3D RICS function; residuals between data and fit are shown above. **G)** Chart showing the diffusion coefficient (µm^2^/s) of TEAD1 and YAP in MCF10A cell nuclei as determine by fluorescence fluctuation spectroscopy. Data presented as mean ± SD, p-values obtained using an unpaired t-test; **** p < 0.0001, n = 10 and 11

Next, we used SMT to further investigate the immobile pool of TEAD1 and YAP by acquiring images of cells with a longer acquisition duration of 500 ms (slow tracking, movie S2). With this imaging regime, the rapidly moving mobile molecules blur into the background and are not detected, allowing one to better characterise the dynamics of the immobile fraction of molecules, such as their DNA dwell times. The majority of both TEAD1 and YAP molecules displayed short DNA dwell times of around 0.5-1 seconds, and a decreasing number of molecules displayed longer residence times, extending up to more than a minute (Fig. 1G and fig. S2f). Many transcription factors have been reported to bind DNA on two broad time scales – long and short – which are thought to reflect specific and non-specific DNA binding, respectively. Accordingly, transcription factor DNA binding times can be calculated using a two-component exponential decay model (*34*). In addition, other studies have reported that transcription factor-DNA binding follows a power-law model, where dwell times occur along a time continuum and are dominated by short binding events (*36*) To test this, we first imaged Halotagged Histone H2B with slow SMT to determine the bleach rate, given that histones can associate with DNA for extended periods (fig. S2e). We then transformed our SMT data by adjusting for photobleaching and assessed whether the data best fit a two-component or three-component exponential models, or a power-law model, using Bayesian Information Criterion (BIC), which takes into account model complexity and goodness of fit. In general, we found that the DNA binding times reliably fit either the power-law model or a two-component exponential model, so we reported both, in addition to the raw data distribution (Fig. 1G and fig. S2, f to i). The power-law exponent for TEAD1 was 0.619 ± 0.011, and for YAP it was 0.734 ± 0.017 (Fig. 1H), indicating that TEAD1 makes more stable contacts with DNA. Consistent with this, bi-exponential fitting indicated that TEAD1 long- and short-DNA dwell times were twice as long as YAP (fig. S2h). This suggests that YAP binds to TEAD molecules that are already DNA associated, or that YAP unbinds TEADs prior to their release from DNA, or a combination of these.

To investigate nuclear TEAD1 and YAP mobility with an independent technique, we employed a novel approach to fluorescence fluctuation spectroscopy termed raster image correlation spectroscopy (RICS). RICS allows extraction of intracellular protein mobility from spatial correlation of fluctuations in fluorescence intensity due to fluorescently tagged protein movement throughout confocal laser scanning microscopy time series data acquisition (*38*). In contrast to our SMT studies, molecular mobility via this approach was defined by a single component, which represents an ensemble of all the detected molecular behaviours. Consistent with our SMT studies, YAP was substantially more mobile than TEAD1 (Fig. 1, I to K). Interestingly, the diffusion coefficients of HaloTagged TEAD1 and YAP (1 μm^2^/s and 3.6 μm^2^/s, respectively) (Fig. 1K) were very similar to our previous observations of endogenously tagged versions of the *Drosophila* orthologues of these proteins in vivo (Scalloped: 1.2 μm^2^/s and Yorkie: 2.9 μm^2^/s) (*39*).

### TEAD1’s ability to bind DNA, not YAP, is the primary determinant of its nuclear behaviour

Most proteins move within cells via diffusion, and this can be influenced by physical interactions they make with other proteins or cellular constituents such as nucleic acids (*40*). Given that TEAD1 can bind both DNA via its TEA domain as well as to proteins like YAP and TAZ, we investigated the relative contributions that these make to its nuclear behaviour. To do this we expressed HaloTagged versions of either wild-type TEAD1, TEAD1 that lacks the TEA domain (TEAD1^ΔDBD^), or TEAD1^Y421H^ (YAP and TEAD1 interact predominantly via a hydrophobic bond between a single amino acid pair (TEAD1 Y421 – YAP S94) and mutation of either of these residues compromises their ability to promote transcription) (*9, 41*). The importance of the YAP-TEAD interaction is further emphasized by the fact that the human genetic disease Sveinnson’s chorioretinal atrophy is caused by the TEAD1^Y421H^ point mutation (*42*). Using fast SMT, we found that TEAD1^ΔDBD^ was substantially more mobile than either TEAD1 or TEAD1^Y421H^, which displayed very similar mobilities (Fig. 2, A to G, and movie S3). This indicates that DNA binding, but not YAP/TAZ binding, is a major determinant of the nuclear mobility of TEAD1. Consistent with published studies, TEAD1’s DNA binding domain also promoted its nuclear localisation, whilst YAP/TAZ binding did not (Fig. 2, A and B). We then used slow SMT to examine the immobile fractions of these different TEAD1 variants. The power-law exponent for overexpressed TEAD1 was 0.739 ± 0.014, and for TEAD1^ΔDBD^ it was 1.089 ± 0.046 (Fig. 2I), indicating that TEAD1 makes far more stable contacts with DNA than TEAD1^ΔDBD^, as expected. The TEAD1^Y421H^ power-law exponent was 0.939 ± 0.029, indicating that when TEAD1 is incapable of binding to YAP/TAZ, it binds to DNA less stably (Fig. 2I). Consistent with this, bi-exponential fitting indicated that long- and short-DNA dwell times were reduced in both the TEAD1^ΔDBD^ and TEAD1^Y421H^ variants (fig. S3, a to d).

**Figure 2.**
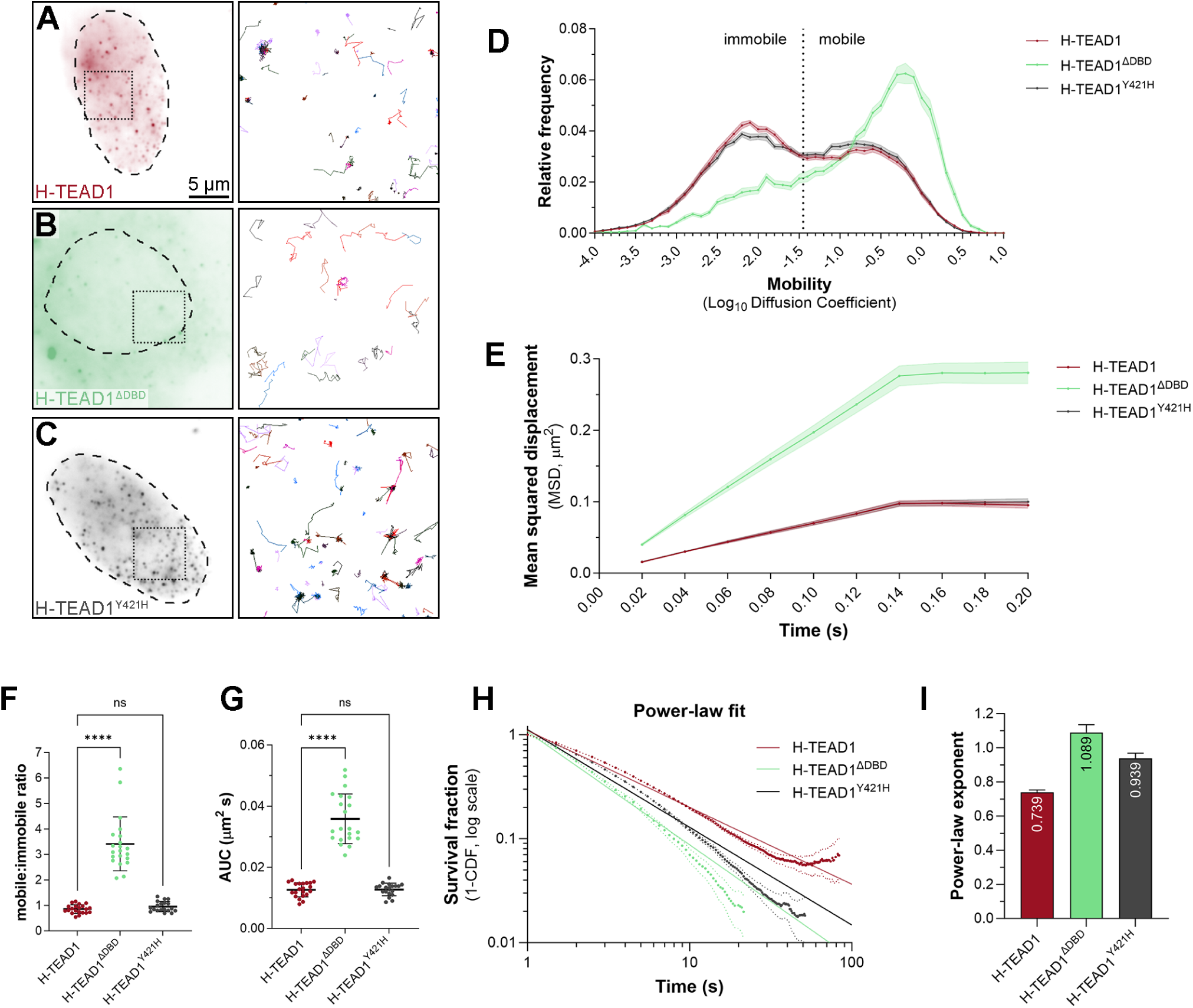
TEAD1’s ability to bind DNA, not YAP, is the primary determinant of its nuclear behaviour. **A-C)** Left panels are average intensity projection images of single HeLa cells expressing Halo-tagged TEAD1, either wild-type (A), protein lacking the TEA DNA binding domain (B), or TEAD where residue 417 is mutated from Y to H (C), imaged at single molecule resolution by HILO. Dashed outlines indicate the nucleus. Right panels show the trajectories of individual TEAD1 molecules tracked over time. Boxed regions in cell images indicate the area corresponding to trajectories. Scale bar is indicated. **D**) Chart showing the relative frequency of molecule mobilities (Log_10_ diffusion coefficient) of TEAD1 (red), TEAD1^ΔDBD^ (green), and TEAD1^Y421H^ (grey) molecules in HeLa cell nuclei over time. Data presented as the mean ± SEM. Mobile and immobile fractions are indicated. Data from 73,348 trajectories from 21 cells for TEAD1, 17,369 trajectories from 21 cells for TEAD1^ΔDBD^, and 67,321 trajectories from 19 cells for TEAD1^Y421H^. **E**) Chart showing the mean squared displacement (µm^2^) of TEAD1 (wild-type and mutant) molecules in HeLa cell nuclei over time. Data presented as the mean ± SEM. **F and G)** Charts showing the mobile to immobile ratio (F) or the area under the curve (G, µm^2^/s) of TEAD1, TEAD1^ΔDBD^, or TEAD1^Y421H^ in HeLa cell nuclei. Data presented as mean ± SD, p values were obtained using a Brown-Forsythe and Welch ANOVA with Dunnett’s T3 multiple comparison test; **** p < 0.0001, ns: not significant, n = 21 cells for TEAD1, 21 cells for TEAD1^ΔDBD^, and 19 cells for TEAD1^Y421H^. **H)** Chart showing the photobleach-corrected survival distribution of TEAD1 (wild-type or mutant) molecules in the nucleus of HeLa cells, with power-law fits (solid lines). Dotted lines are 99% CI. N = 202,613 trajectories from 26 cells for TEAD1, 38,766 trajectories from 22 cells for TEAD1^ΔDBD^, and 83,778 trajectories from 21 cells for TEAD1^Y421H^. **I)** Bar chart showing the power-law exponents of TEAD1 proteins. Error bars indicate 95% CI.

### TEADs have a major influence on YAP’s nuclear behaviour

We next examined how TEADs impact YAP nuclear behaviour and chromatin association by studying a mutant YAP protein (YAP-S94A, which cannot bind to TEADs) (*9*), as well as the impact of TEAD1 overexpression on YAP. Fast SMT revealed that TEADs have a major influence on YAP behaviour; while YAP exhibited both mobile and immobile behaviour in the nucleus, YAP^S94A^ was almost entirely mobile (Fig. 3, A to G). RICS also showed that nuclear YAP^S94A^ was far more mobile than YAP (fig. S4a). In addition, TEAD1 overexpression significantly decreased the overall nuclear mobility of YAP by SMT (Fig. 3, A to G), and transformed it’s overall mobility distribution to almost match TEAD1 (compare to Figs. 1 and 2). These phenotypes were evident with every quantitative analysis we performed, i.e. mobility, mean squared displacement, area under the curve and mobile to immobile ratio (Fig. 3, D to G).

**Figure 3.**
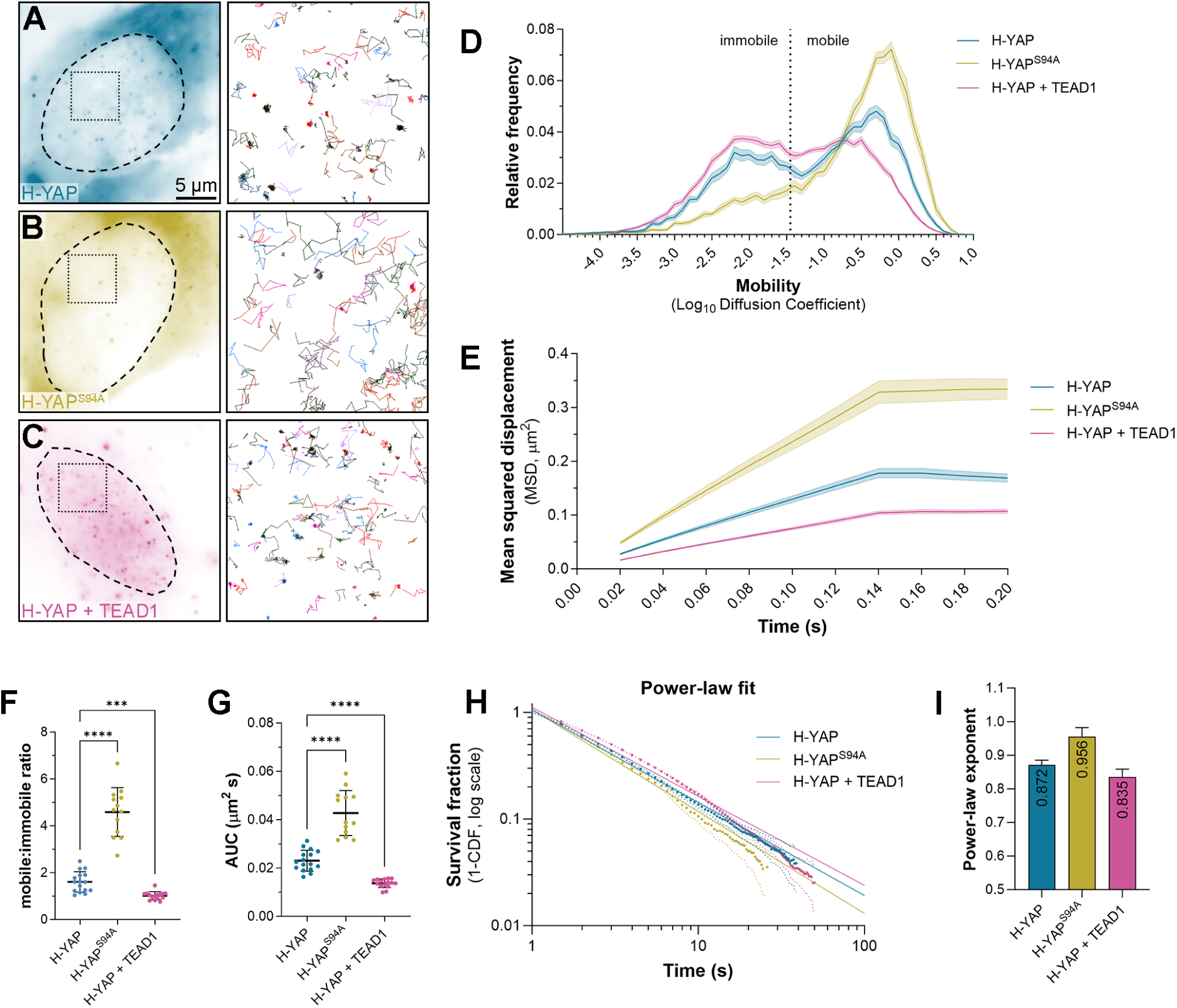
TEADs have a major influence on YAP’s nuclear behaviour. **A-C)** Left panels are average intensity projection images of single HeLa cells expressing Halo-tagged YAP, either wild-type (A), YAP where residue 94 is mutated from S to A (B), or wild-type YAP in the presence of overexpressed TEAD1 (C), imaged at single molecule resolution by HILO. Dashed outlines indicate the nucleus. Right panels show the trajectories of individual YAP molecules tracked over time. Boxed regions in cell images indicate the area corresponding to trajectories. Scale bar is indicated. **D**) Chart showing the relative frequency of molecule mobilities (Log_10_ diffusion coefficient) of YAP (blue), YAP^S94A^ (olive), and YAP with overexpressed TEAD1 (pink) molecules in HeLa cell nuclei over time. Data presented as the mean ± SEM. Mobile and immobile fractions are indicated. Data from 14,648 trajectories from 16 cells for YAP, 13,785 trajectories from 13 cells for YAP^S94A^, and 52,442 trajectories from 15 cells for YAP with overexpressed TEAD1. **E**) Chart showing the mean squared displacement (µm^2^) of YAP (wild-type and mutant) molecules in HeLa cell nuclei over time. Data presented as the mean ± SEM. **F and G)** Charts showing the mobile to immobile ratio (F) or the area under the curve (G, µm^2^/s) of YAP, YAP^S94A^, or YAP with overexpressed TEAD1 in HeLa cell nuclei. Data presented as mean ± SD, p values were obtained using a Brown-Forsythe and Welch ANOVA with Dunnett’s T3 multiple comparison test; **** p < 0.0001, *** p < 0.001, ns: not significant, n = 16 cells for YAP, 13 cells for YAP^S94A^, and 15 cells for YAP with overexpressed TEAD1. **H)** Chart showing the photobleach-corrected survival distribution of YAP (wild-type or mutant) molecules in the nucleus of HeLa cells, with power-law fits (solid lines). Dotted lines are 99% CI. N = 62,126 trajectories from 24 cells for YAP, 33,571 trajectories from 13 cells for YAP^S94A^, and 126,127 trajectories from 16 cells for YAP with overexpressed TEAD1. **I)** Bar chart showing the power-law exponents of TEAD1 proteins. Error bars indicate 95% CI.

Next, we examined the impact of TEADs on YAP chromatin association, using slow SMT. The power-law exponent for YAP was 0.872 ± 0.014 (Fig. 3, H and I), whilst for YAP^S94A^ it was 0.956 ± 0.026, indicating less stable DNA binding. In comparison to YAP alone, overexpression of TEAD1 slightly reduced YAP’s power-law exponent to 0.835 ± 0.023. Two-component fitting of the DNA dwell times showed that YAP^S94A^ had reduced short and long dwell times relative to YAP, but overexpression of TEAD1 had no major impact on YAP’s dwell times; rather, it increased the fraction of longer-lived dwell times from ∼15% to 20% (fig. S4, b to e). Taken together with the fast-tracking results, these data show that TEADs are a major regulator of YAP’s nuclear dynamics, and, consistent with numerous previous studies, facilitates YAP-DNA binding.

### Hippo signalling limits the DNA dwell times of YAP and TEAD1

How signalling pathways impact the biophysical behaviour and DNA binding of their downstream transcription factors has only begun to be explored. Exceptions include the Notch and Glucocorticoid pathways, which both stimulate transcription by inducing an increase in dwell time of their relative transcription factors (*43, 44*). The best characterised way in which Hippo signalling controls transcription is by reducing the nuclear pool of YAP by phosphorylating it (*1–4*). To study the impact of Hippo signalling on YAP and TEAD1 nuclear behaviours and DNA binding we used recently developed LATS inhibitors (LATSi), which rapidly suppress LATS1/2-dependent phosphorylation of YAP (*45*). MCF10A cells were treated with LATSi or DMSO for 2 hours and YAP and TEAD1 imaged by single molecule tracking, as we observed robust loss of YAP-S127 phosphorylation (fig. S5a). Consistent with previous studies that assessed YAP localisation at the bulk protein level by immunofluorescence, we observed a substantial increase in nuclear Halo-YAP following LATSi treatment, while the localisation of TEAD1 was unaffected (Fig. 4, A to C). Fast SMT revealed that TEAD1 nuclear mobility was decreased slightly following LATS1/2 inhibition (Fig. 4D). By contrast, YAP mobility was slightly increased by LATS1/2 inhibition (Fig. 4E). These changes were evident when assessing mean squared displacement and area under the curve (Fig. 4, F and H). A significant change was also observed in the mobile to immobile ratio of TEAD1; for YAP this trended towards significance (p=0.169) (Fig. 4G). When we performed the same experiments using RICS, we were unable to detect any significant differences in YAP or TEAD1 mobility when LATS activity was inhibited (fig. S5b).

**Figure 4.**
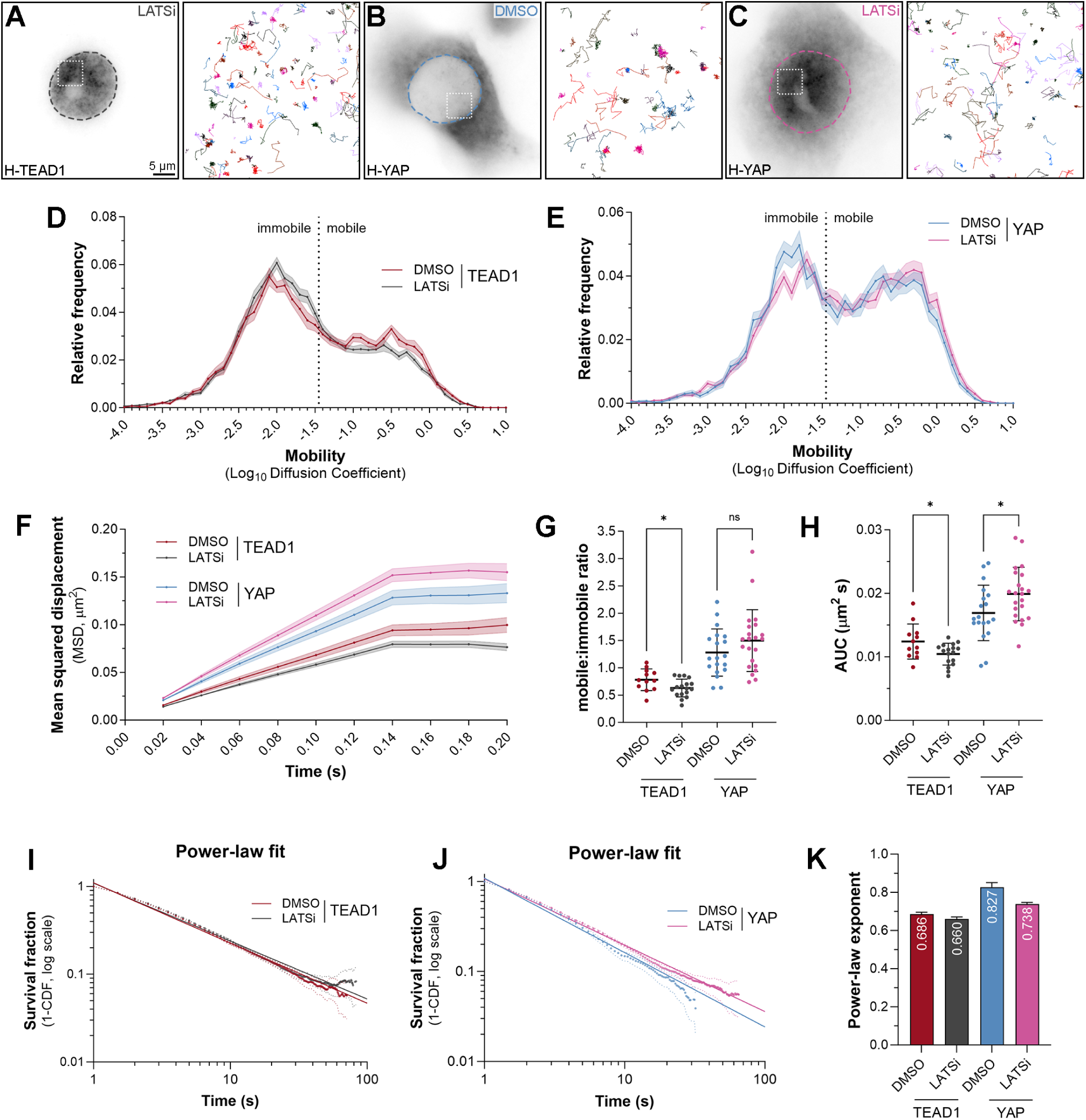
Hippo signalling limits the DNA dwell times of YAP and TEAD1. **A-C)** Left panels are images of single MCF10A cells expressing Halo-tagged TEAD1 (A) or YAP (B and C), imaged with widefield epifluorescence. The cells on the left (A) and right (C) were treated with 3 µM LATSi for 2 hours, whilst the central cell was treated with DMSO. Dashed outlines indicate the nucleus. Right panels show the trajectories of individual TEAD1 molecules tracked over time. Boxed regions in cell images indicate the area corresponding to trajectories. Scale bar is indicated. **D and E)** Chart showing the relative frequency of molecule mobilities (Log_10_ diffusion coefficient) of TEAD1 (D) or YAP (E) in in MCF10A cell nuclei over time. Cells were treated with DMSO or LATSi. Data presented as the mean ± SEM. Mobile and immobile fractions are indicated. Data from 14,053 trajectories from 12 cells for TEAD1 (DMSO), 20,093 trajectories from 17 cells for TEAD1 (LATSi), 6,386 trajectories from 19 cells for YAP (DMSO), and 10,752 trajectories from 22 cells for YAP (LATSi). **F**) Chart showing the mean squared displacement (µm^2^) of TEAD1 or YAP molecules in MCF10A cell nuclei over time. Data presented as the mean ± SEM. Cells were treated with DMSO or LATSi. **G and H)** Charts showing the mobile to immobile ratio (G) or the area under the curve (H, µm^2^/s) of TEAD1 and YAP molecules in MCF10A cell nuclei. Cells were treated with DMSO or LATSi. Data presented as mean ± SD, p-values obtained using an unpaired t-test; * p < 0.05, ns: not significant, n = 12 and 17 cells for TEAD1 and n = 19 and 22 cells for YAP (DMSO and LATSi, respectively). **I and J)** Chart showing the photobleach-corrected survival distribution of single Halo-TEAD1 or Halo-YAP molecules in the nucleus of MCF10A cells, with power-law fits (solid lines). Cells were treated with DMSO or LATSi. Dotted lines are 99% CI. n = 54,886 trajectories from 21 cells for TEAD1 (DMSO), 68,992 trajectories from 17 cells for TEAD1 (LATSi), 14,104 trajectories from 19 cells for YAP (DMSO), and 46,056 trajectories from 22 cells for YAP (LATSi). **K)** Bar chart showing the power-law exponents of YAP and TEAD1 in MCF10A cells treated with either DMSO or LATSi. Error bars indicate 95% CI.

We next examined whether blocking Hippo signalling affected YAP and TEAD1 chromatin association, by performing slow SMT in MCF10A cells treated with LATSi for 2 hours. Interestingly, the power-law exponents of both and YAP and TEAD1 were lower in LATSi-treated cells, indicating that their DNA dwell times were extended (Fig. 4, I to K). We then extracted long and short DNA dwell times from the two-component exponential fit (fig. S5, d to f). These revealed that both YAP and TEAD1 long-dwell times doubled after LATSi treatment: YAP increased from ∼18.3 to ∼41.4 seconds, whilst TEAD1 went from ∼42.2 to ∼82.5 seconds. Similarly, short-lived dwell times were also slightly increased. In contrast, both TEAD1 and YAP had a slightly reduced fraction of long-lived binding events (fig. S5g). Therefore, Hippo signalling normally limits the amount of nuclear YAP, and the time that both YAP and TEAD1 associate with the genome.

### YAP and TEAD1 associate with DNA for extended periods in cells with intrinsically low Hippo signalling

To investigate the impact of Hippo signalling in an independent fashion and a more physiological setting, we leveraged knowledge of Hippo signalling in the early mammalian embryo, where it is essential for the very first cell fate choice, i.e., trophectoderm versus inner cell mass (*46*). Hippo signalling is high in the inner cell mass, which goes on to develop the embryo, whilst it is low in the trophectoderm, which forms the placenta (*46*). Differential Hippo signalling in these two juxtaposed cell types results in YAP/TEAD activity in the trophectoderm and low YAP/TEAD activity in the inner cell mass (*46*). As such, it is an ideal biological system to explore YAP/TEAD dynamics.

However, the physical dimensions of the early mammalian embryo were not compatible for reliable single molecule imaging as HILO imaging significantly deteriorates the further the sample is from the coverslip. Therefore, to image YAP and TEAD1 at single molecule resolution in living cells resembling the early embryo, we used two well-known 2D cellular models; induced pluripotent stem cells (iPSC), to model the epiblast, and induced trophoblast stem cells (iTSC), to model the trophoblasts (*47, 48*). To do this we stably expressed Dox-inducible Halo-YAP and Halo-TEAD1 in iPSCs or iTSCs. Both YAP and TEAD1 were nuclear in both cell types when grown in monocultures, as determined with YAP antibodies (fig. S6a). However, when these cells were grown as iTSC/iPSC co-cultures, iPSC’s grew together in spheres, where YAP was cytoplasmic, as it is in the inner cell mass of the early embryo (fig. S6b and d). By contrast, YAP remained nuclear in iTSC’s that surround the iPSC islands, as it is in the trophectoderm (fig. S6b and c). As such, this iTSC/iPSC co-culture system provided the ideal setting to study YAP and TEAD1 behaviour at single molecule resolution in cells with intrinsic differences in Hippo signalling.

Fast SMT revealed that nuclear TEAD1 dynamics were indistinguishable in both iPSC and iTSC (Fig. 5, D to H). Even though YAP was more nuclear iTSC’s than iPSC’s when grown in co-culture, YAP’s nuclear dynamics were also very similar in both cell types (Fig. 5E). YAP mobility, as assessed by mean squared displacement and area under the curve did not differ significantly, whilst the mobile to ratio immobile ratio of YAP was slightly lower in iTSC (Fig. 5, F to H). Next, we assessed the DNA association dynamics of TEAD1 and YAP in these two cell types, using slow SMT. The TEAD1 power-law exponent was 0.642 ± 0.008 in iPSC and 0.529 ± 0.005 in iTSC, indicating that TEAD1 binds DNA substantially longer in iTSC than iPSC (Fig. 5, I to K). This was also evidenced when fitting the data to a two-component exponential decay models, where the long binding time of TEAD1 increased from ∼50 seconds to ∼90.9 seconds (fig. S7, b and d). Similar results were observed for YAP, which had a power-law exponent of 0.723 ± 0.014 in iPSC and 0.653 ± 0.007 in iTSC (Fig. 5, J and I). When fitted to a two-component exponential decay model, the YAP long binding time increased from ∼20.3 seconds to ∼40.9 seconds (fig. S7, c and d). Slight increases were also observed for both YAP and TEAD1 short dwell times, and minor changes in the long and short binding fractions (fig. S7, d and e). Therefore, both TEAD1 and YAP bind chromatin for longer in cells with lower intrinsic Hippo pathway activity, and where YAP is more nuclear.

**Figure 5.**
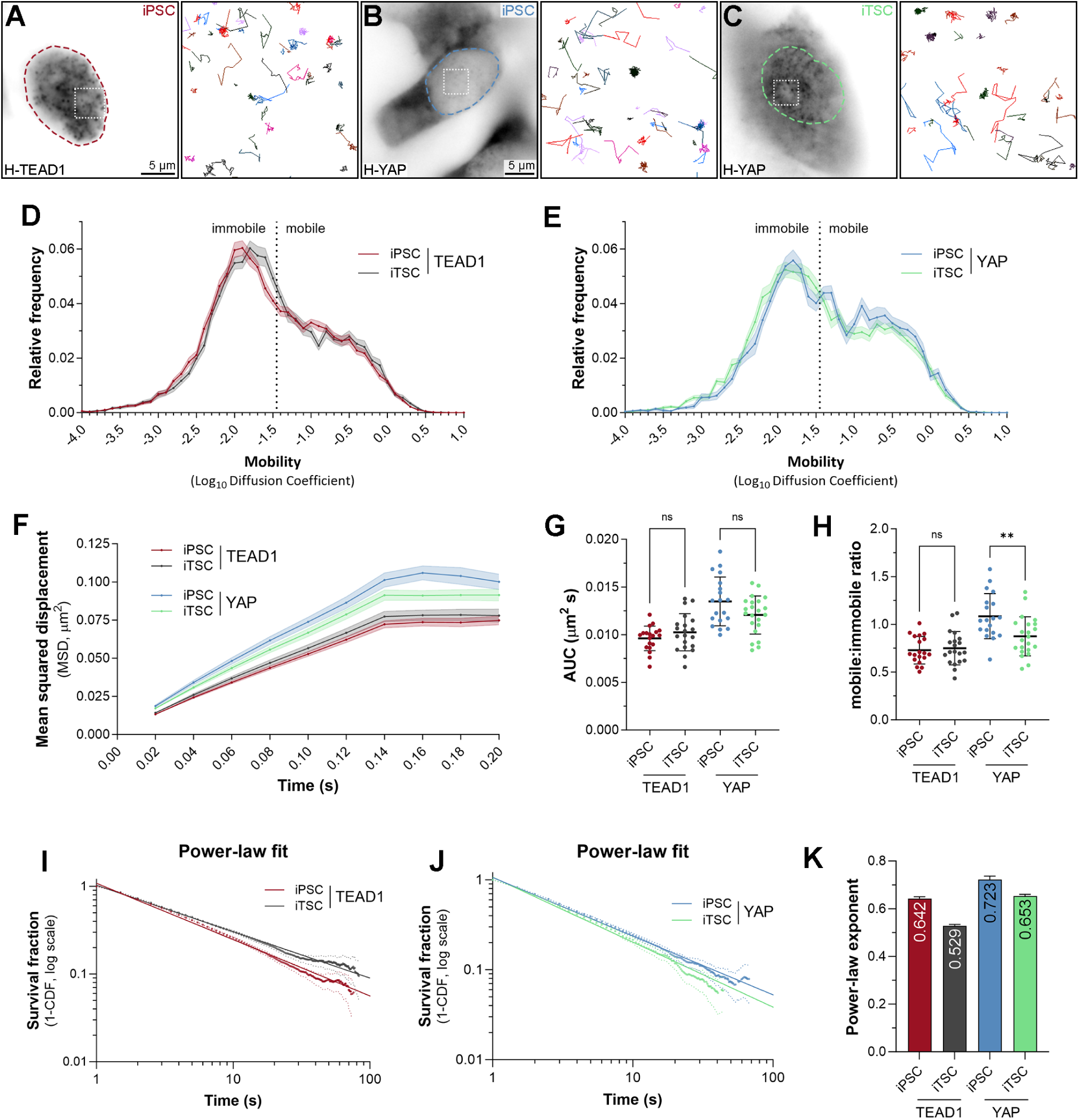
Longer YAP and TEAD1 DNA dwell times in cells with intrinsically high YAP/TEAD activity. **A-C)** Left panels are images of single iPS or iTS cells expressing Halo-tagged TEAD1 (A) or YAP (B and C), imaged with widefield epifluorescence. Cells are from co-cultures with wild-type iPS or iTS cells. Dashed outlines indicate the nucleus. Right panels show the trajectories of individual TEAD1 or YAP molecules tracked over time. Boxed regions in cell images indicate the area corresponding to trajectories. Scale bars are indicated. **D and E)** Chart showing the relative frequency of molecule mobilities (Log_10_ diffusion coefficient) of TEAD1 (D) or YAP (E) in iPS or iTS cell nuclei. Data presented as the mean ± SEM. Mobile and immobile fractions are indicated. Data from 17,036 trajectories from 19 cells for TEAD1 (iPS cells), 17,257 trajectories from 20 cells for TEAD1 (iTS cells), 6,874 trajectories from 19 cells for YAP (iPS cells), and 23,382 trajectories from 22 cells for YAP (iTS cells). **F)** Chart showing the mean squared displacement (µm^2^) of TEAD1 or YAP molecules in iPS or iTS cell nuclei over time. Data presented as the mean ± SEM. **G and H)** Charts showing the mobile to immobile ratio (G) or the area under the curve (H, µm^2^/s) of TEAD1 and YAP molecules in iPS or iTS cell nuclei. Data presented as mean ± SD, p-values obtained using an unpaired t-test; ** p < 0.01, ns: not significant, n = 19 and 20 cells for TEAD1 and n = 19 and 22 cells for YAP (iPS and iTS cells, respectively). **I and J)** Chart showing the photobleach-corrected survival distribution of single Halo-TEAD1 or Halo-YAP molecules in the nucleus of co-cultured iPS or iTS cells, with power-law fits (solid lines). Dotted lines are 99% CI. n = 47,191 trajectories from 22 cells for TEAD1 (iPS cells), 42,339 trajectories from 22 cells for TEAD1 (iTS cells), 13,381 trajectories from 25 cells for YAP (iPS cells), and 26,931 trajectories from 23 cells for YAP (iTS cells). **K)** Bar chart showing the power-law exponents of YAP and TEAD1 in co-cultured iPS and iTS cell nuclei. Error bars indicate 95% CI.

### TEAD1 mobility is impaired in nuclear condensates

Throughout the course of imaging we noted that DNA binding times occurred on a broad continuum. Therefore, we explored whether it was heterogeneous across the nucleus, particularly given reports that both YAP and TAZ form nuclear condensates (*26–29*), which are membraneless subdomains where these proteins are found in higher concentrations. YAP and TAZ condensates are also enriched for TEAD transcription factors and chromatin-regulatory proteins like Brd4, that promote transcription (*26–29*). We also observed these condensates in HeLa cells expressing both YAP-GFP and Halo-TEAD1 (movie S4 and Fig. 6G). YAP exhibits reduced mobility in nuclear condensates (*27*) and although TEAD is present in these condensates, its behaviour in these regions has not been addressed. To investigate this, we performed SMT in Halo-TEAD1 transduced MCF10A cells, without overexpressing YAP, and quantified TEAD1 behaviour in condensates compared with non-punctate regions of nuclei (fig. S8a). In condensates, TEAD1 mobility was substantially impaired, as determined by fast SMT (Fig. 6, A to D). To further address whether the observed continuum of TEAD1’s DNA binding durations are affected by sub-nuclear domains, we performed slow SMT. We found that TEAD1’s DNA binding behaviour best fit a power-law model in both condensates and non-punctate nuclear regions (Fig. 6H). The TEAD1 power-law exponent was 0.647 ± 0.055 in condensates, compared to 0.762 ± 0.038 in non-punctate regions, indicating that TEAD1 dwell times were extended in condensates (Fig. 6I). Bi-exponential models, on the other hand, indicated that the fraction of long-lived DNA binding events was greater in condensates, with ∼35% of tracks being long, relative to ∼23% in non-puncta (fig. S8c and d to g).

**Figure 6.**
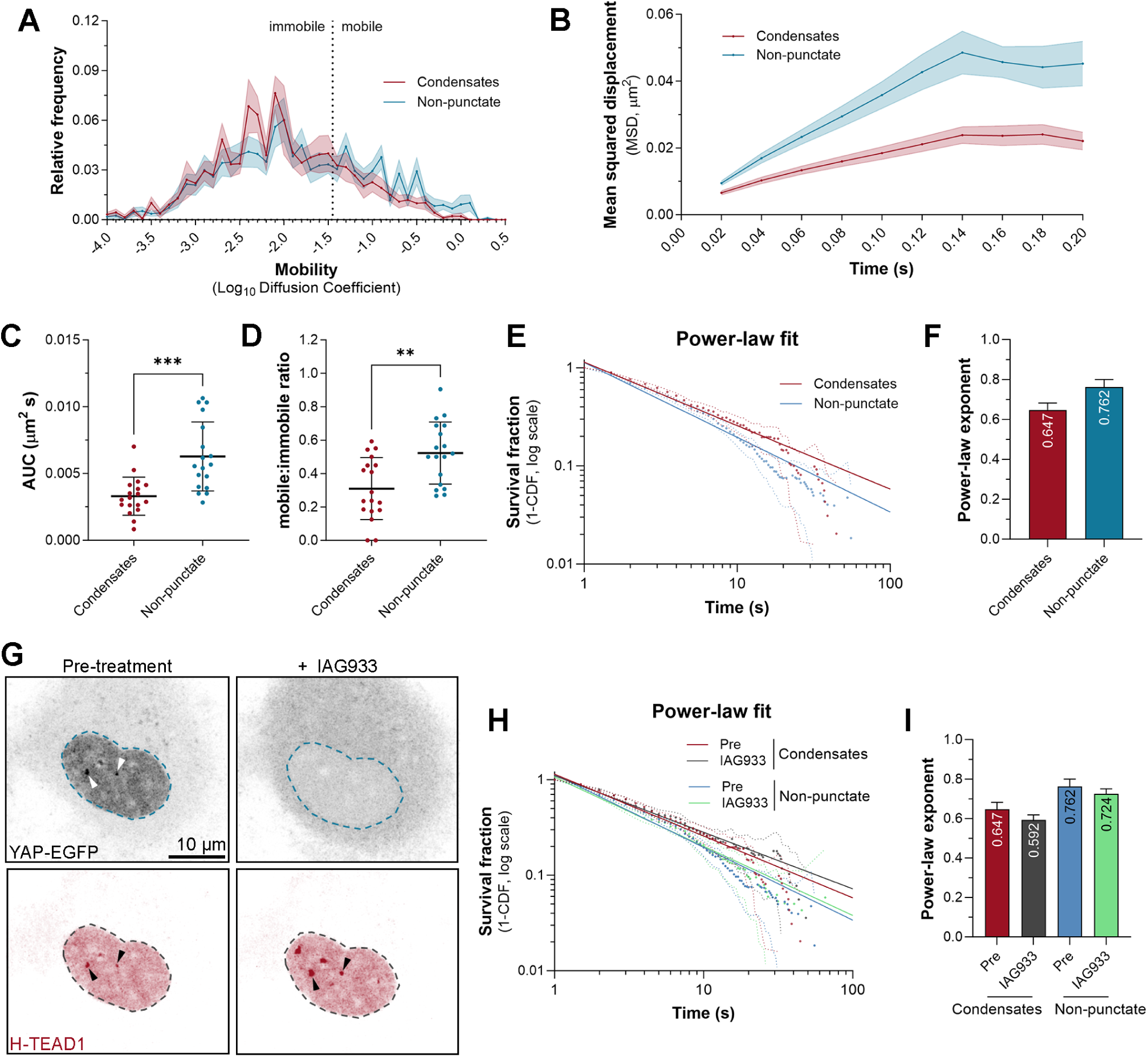
TEAD1 exhibits reduced mobility and extended DNA binding behaviour in nuclear condensates. **A**) Chart showing the relative frequency of molecule mobilities (Log_10_ diffusion coefficient) of Halo-tagged TEAD1 in MCF10A cell nuclear condensates and nuclear regions with no visible condensates. Data presented as the mean ± SEM. Mobile and immobile fractions are indicated. Data from 1,247 trajectories in condensates, and 955 trajectories in non-punctate nuclear regions, from 18 cells. **B**) Chart showing the mean squared displacement (µm^2^) of TEAD1 molecules in MCF10A cell nuclear condensates and non-punctate nuclear regions, over time. Data presented as the mean ± SEM. **C and D)** Charts showing the mobile to immobile ratio (C) or the area under the curve (D, µm^2^/s) of TEAD1 molecules in MCF10A cell nuclear condensates and non-punctate nuclear regions. Data presented as mean ± SD, p-values obtained using an unpaired t-test; *** p < 0.001, ** p < 0.01, n =18 cells. **E)** Chart showing the photobleach-corrected survival distribution of Halo-TEAD1 molecules in MCF10A cell nuclear condensates and non-punctate nuclear regions, with power-law fits (solid lines). Dotted lines are 99% CI. n = 1,080 trajectories in condensates, and 2,628 trajectories in non-punctate regions, from 14 cells. **F)** Bar chart showing the power-law exponents of TEAD1 in MCF10A cell nuclear condensates and nuclear regions with no visible condensates. Error bars indicate 95% CI. **G)** Maximum intensity projection confocal microscope images of single HeLa cells expressing YAP-EGFP (black) and Halo-tagged TEAD1 (red) pre-and post-treatment with 3 μM IAG933 for ∼8 minutes. Dashed outlines indicate the nucleus. Black arrows indicate TEAD1 condensates, and white arrows indicate YAP condensates in the pre-treatment cells. Scale bars are indicated. **H)** Chart showing the photobleach-corrected survival distribution of Halo-TEAD1 molecules in MCF10A cell nuclear condensates and non-punctate nuclear regions, pre- and post-treatment with 3 μM IAG933 for 30 mins, with power-law fits (solid lines). Dotted lines are 99% CI. Pre-treatment, n = 1,080 trajectories in condensates, and 2,628 trajectories in non-punctate regions, from 14 cells. Post-treatment, n = 672 trajectories in condensates, and 1,465 trajectories in non-punctate regions. **I)** Bar chart showing the power-law exponents of TEAD1 in MCF10A cell nuclear condensates and regions with no visible condensates, pre- and post-treatment with 3 μM IAG933 for 30 mins. Error bars indicate 95% CI.

To investigate the impact of the YAP and TEAD1 interaction on nuclear condensates further, we utilised a newly reported compound that disrupts the YAP/TEAD protein-protein interaction and alters their genome occupancy at target sites (*49*). Initially, we examined the impact of IAG933 on HeLa cells transiently overexpressing YAP-GFP and Halo-TEAD1 at the bulk protein level, using live confocal microscopy. Remarkably, within seconds of treatment of cells with IAG933, most YAP nuclear condensates were no longer visible, and YAP protein began to accumulate in the cytoplasm (Fig. 6G and fig. S8d). In contrast, TEAD1’s nuclear localisation was unaffected and, surprisingly, TEAD1 condensates were unperturbed or increased in size and number (Fig. 6G and fig. S8d). We then performed slow SMT on TEAD1 in nuclear condensates pre- and post- the addition of IAG933 in live MCF10A cells to assess any effects on its DNA binding behaviour.

Power-law models indicated that TEAD1 DNA dwell times were extended following IAG933 treatment, in both condensates and in non-punctate regions (Fig. 6, H and I). Bi-exponential fitting agreed with this impact on condensate-associated TEAD1, as the long and short dwell times were longer following IAG933 treatment (fig. S8, e to g).

### A cancer-associated YAP fusion protein displays different nuclear behaviour than YAP

Multiple Hippo pathway genes are mutated in different human cancers, including YAP and TAZ, which, following chromosomal translocations, can fuse with different transcription factors (*32*). One such fusion protein, YAP-TFE3, occurs in 10-20% of epithelioid hemangioendotheliomas (*33*), a vascular tumour that is caused by fusion of the N-terminus of either YAP or TAZ with TFE3 or CAMTA1, respectively (*32*). To investigate whether the biophysical behaviour of cancer-associated YAP fusion proteins differs from YAP, we performed single molecule tracking of HaloTagged YAP-TFE3. Fast tracking revealed that, like YAP and TEAD1, overexpressed YAP-TFE3 exhibits both mobile and immobile behaviour, but is less mobile than either YAP or TEAD1 (Fig. 7, compare to Figs. 2 and 3). Multiple independent studies have revealed that YAP-TFE3 drives oncogenic transcription at least in part through TEADs. Therefore, to investigate the relative importance of TEAD binding for YAP-TFE3 nuclear behaviour, we imaged a version of YAP- TFE3 that cannot interact with TEADs (Halo-YAP^S94A^-TFE3). YAP^S94A^-TFE3 was substantially more mobile than YAP-TFE3 (Fig. 7, A to D), indicating that TEADs are a major determinant of YAP-TFE3 behaviour, as they are for YAP. This was further confirmed by the finding that TEAD1 overexpression further decreased the mobility of YAP-TFE3 (Fig. 7, A to D). Additionally, TEAD1 mobility was strikingly reduced by YAP-TFE3 overexpression (Fig. 7, G to J). Therefore, these studies reveal that cancer-associated alleles of YAP transform the nuclear dynamics of the wildtype transcription regulatory protein.

**Figure 7.**
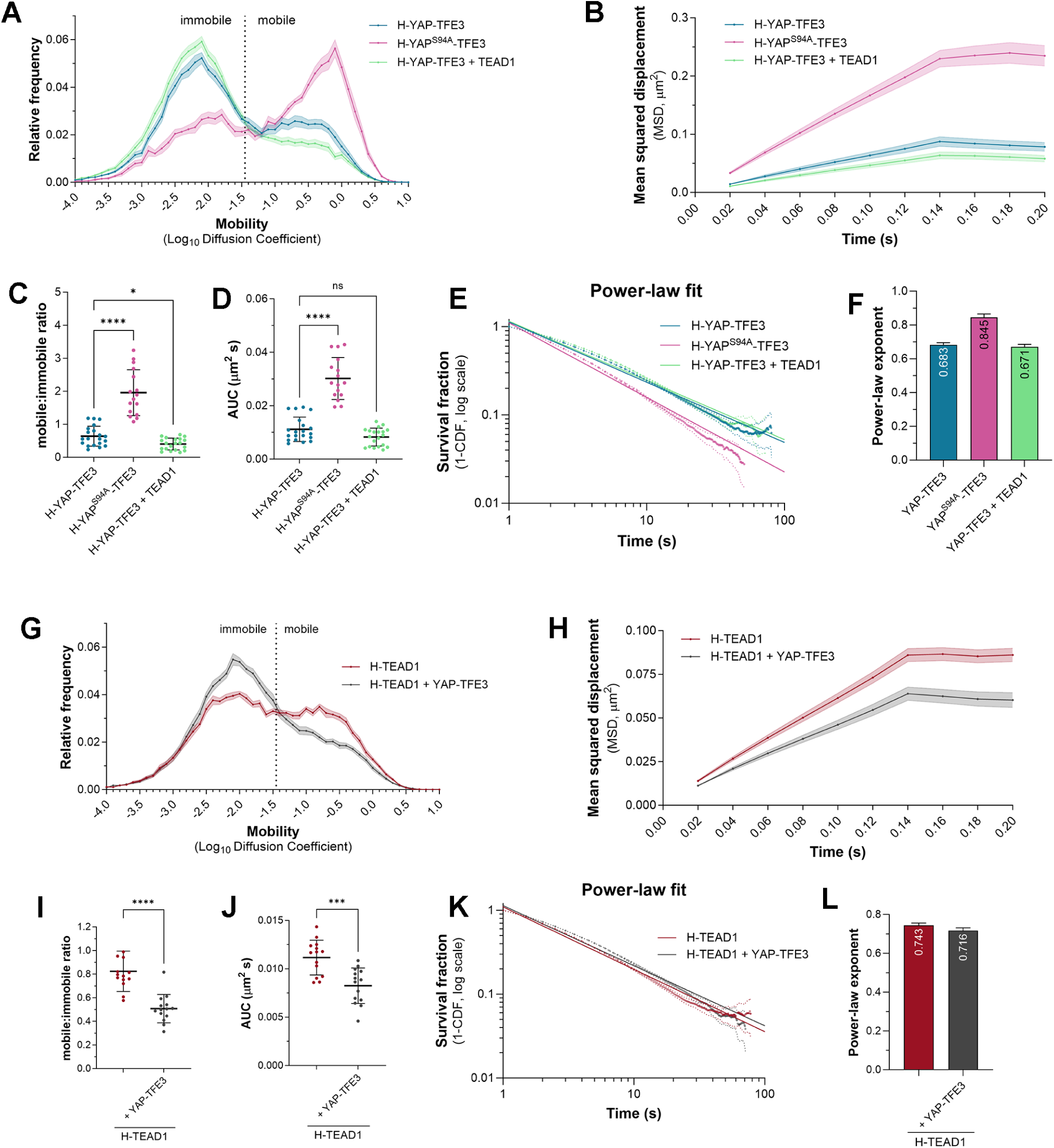
The nuclear behaviour of the YAP-TFE3 oncogenic fusion protein differs substantially from YAP but is still influenced by TEADs. **A**) Chart showing the relative frequency of molecule mobilities (Log_10_ diffusion coefficient) of YAP-TFE3 (blue), YAP^S94A^-TFE3 (pink), and YAP-TFE3 with overexpressed TEAD1 (green) molecules in HeLa cell nuclei over time. Data presented as the mean ± SEM. Mobile and immobile fractions are indicated. Data from 42,829 trajectories from 20 cells for YAP-TFE3, 34,229 trajectories from 16 cells for YAP^S94A^-TFE3, and 71,291 trajectories from 20 cells for YAP-TFE3 with overexpressed TEAD1. **B**) Chart showing the mean squared displacement (µm^2^) of YAP-TFE3, YAP^S94A^-TFE3, and YAP- TFE3 with overexpressed TEAD1, molecules in HeLa cell nuclei over time. Data presented as the mean ± SEM. **C and D)** Charts showing the mobile to immobile ratio (C) or the area under the curve (D, µm^2^/s) of YAP-TFE3, YAP^S94A^-TFE3, or YAP-TFE3 with overexpressed TEAD1, in HeLa cell nuclei. Data presented as mean ± SD, p values were obtained using a Brown-Forsythe and Welch ANOVA with Dunnett’s T3 multiple comparison test; **** p < 0.0001, * p < 0.05, ns: not significant, n = 20 cells for YAP-TFE3, 16 cells for YAP^S94A^-TFE3, and 20 cells for YAP-TFE3 with overexpressed TEAD1. **E**) Chart showing the photobleach-corrected survival distribution of YAP-TFE3, YAP^S94A^-TFE3, or YAP-TFE3 with overexpressed TEAD1 in the nuclei of HeLa cells, with power-law fits (solid lines). Dotted lines are 99% CI. n = 167,283 trajectories from 27 cells for YAP-TFE3, 104,009 trajectories from 19 cells for YAP^S94A^-TFE3, and 86,983 trajectories from 20 cells for YAP-TFE3 with overexpressed TEAD1. **F**) Bar chart showing the power-law exponents of YAP-TFE3, YAP^S94A^-TFE3, or YAP-TFE3 with overexpressed TEAD1, in the nuclei of HeLa cells, with 95% CI. **G**) Chart showing the relative frequency of molecule mobilities (Log_10_ diffusion coefficient) of Halo-tagged TEAD1 alone or in combination with YAP-TFE3-GFP overexpression. Data presented as the mean ± SEM. Mobile and immobile fractions are indicated. Data from 46,400 trajectories from 14 cells for TEAD1 alone, and 50,235 trajectories from 14 cells for TEAD1 with overexpressed YAP-TFE3. **H**) Chart showing the mean squared displacement (µm^2^) of TEAD1 molecules in the presence of absence of overexpressed YAP-TFE3-GFP in HeLa cell nuclei over time. Data presented as the mean ± SEM. **I and J)** Charts showing the mobile to immobile ratio (I) or the area under the curve (J, µm^2^/s) of TEAD1 molecules in the presence or absence of overexpressed YAP-TFE3-GFP in HeLa cell nuclei. Data presented as mean ± SD, p values were obtained using a Brown-Forsythe and Welch ANOVA with Dunnett’s T3 multiple comparison test; **** p < 0.0001, *** p < 0.001, n = 14 cells for TEAD1 with and without overexpressed YAP-TFE3-GFP. **K)** Chart showing the photobleach-corrected survival distribution of Halo-TEAD1 molecules in nuclei of HeLa cells in the presence or absence of YAP-TFE3-GFP, with power-law fits (solid lines). Dotted lines are 99% CI. n = 121,486 trajectories from 16 cells for TEAD1 alone, and 102,439 trajectories from 15 cells for TEAD1 with overexpressed YAP-TFE3-GFP. **K)** Bar chart showing the power-law exponents of TEAD1 molecules in nuclei of HeLa cells in the presence or absence of YAP-TFE3-GFP, with 95% CI.

To investigate YAP-TFE3 DNA binding, we performed slow tracking. The DNA dwell time of YAP-TFE3 was longer than overexpressed YAP or TEAD1, as revealed by a power law exponent of 0.683 ± 0.012 (Fig. 7F, compare to Fig. 3I and 7J). TEADs also had an impact on YAP-TFE3 DNA binding; YAP^S94A^-TFE3 had a power-law exponent of 0.845 ± 0.019, indicating that association with TEADs facilitate more stable DNA contacts. However, TEAD1 overexpression did not significantly alter YAP-TFE3’s power-law exponent (0.671 ± 0.015; Fig. 7F). Consistent with the finding that TEAD’s influence YAP-TFE3’s DNA residence times, bi-exponential fitting showed that YAP^S94A^-TFE3’s long- and short- dwell times were reduced by more than half relative to YAP-TFE3 (fig. S9b and c). Notably, preventing the ability of YAP-TFE3 to bind TEADs did not cause the prevent all immobile behaviour of YAP-TFE3, presumably because TFE3 also has a DNA binding domain, and we previously found using genomics experiments that YAP-TFE3 binds the genome via both TEADs and TFE3 (*50*). Despite TEAD1 becoming less mobile in the presence of YAP-TFE3, slow tracking indicated no substantial changes to its DNA dwell times, as determined by the power-law fit (Fig. 7, K and L).

## DISCUSSION

Using advanced microscopy techniques, we revealed new insights into the molecular mobility and DNA binding properties of the key transcription pair, YAP and TEADs, and how these features are regulated by Hippo pathway signalling. We found that both YAP and TEAD1 exhibit mobile and immobile behaviour in the nucleus, with YAP being more mobile than TEAD1. YAP’s mobility was greatly impacted by its ability to form a physical complex with TEADs, whilst YAP only had a minor influence on TEAD1 mobility. Both proteins bound chromatin across broad timescales from fractions of seconds through to minutes, with YAP generally having shorter DNA dwell times than TEAD1, which suggests three possibilities: 1) TEAD reside on DNA and YAP binds it there, and they unbind DNA simultaneously; 2) YAP and TEAD first form a heterodimer in the nucleoplasm, bind DNA together and then YAP disengages from TEAD before TEAD unbinds DNA; or 3) a combination of these two possibilities. Our finding that YAP diffuses throughout the nucleus approximately three times faster than TEAD1 argues in favour of the first possibility, i.e., that YAP and TEAD1 predominantly form a heterodimer on DNA, rather than diffusing throughout the nucleus as a pre-formed heterodimer. This is further supported by the recent finding that the *Drosophila* orthologues of YAP and TEADs – Yorkie and Scalloped – diffuse at almost the same rates in nuclei of *Drosophila* tissues as YAP and TEAD1 (*39*).

Single molecule tracking microscopy also revealed an important new mechanism by which the Hippo pathway controls transcription, i.e. Hippo signalling normally limits the time that both YAP and TEAD1 bind to chromatin. Coupled with recent observations in *Drosophila* where overexpression of the Yorkie co-activator extended the DNA dwell times of Scalloped (*39*), this suggests that in addition to regulating the nucleocytoplasmic distribution of YAP, the Hippo pathway controls transcription by regulating YAP/TEAD chromatin dwell time. The fact that similar discoveries have been made in both *Drosophila* and human cells, further underscores how key features of this ancient signalling pathway have been conserved throughout evolution (*51, 52*). We made these observations by acutely perturbing Hippo signalling using chemical inhibitors of the LATS1/2 kinases, as well as with a cell co-culture model that recapitulates key features of the pre- implantation mammalian embryo, i.e. that Hippo signalling is high in the inner cell mass, and low in the trophectoderm. In recent years, multiple groups have developed in vitro models of the human blastocyst, which form three-dimensional embryo-like structures (*47, 53*). Despite their utility for studying features of early human life, the three-dimensional structures they form present challenges for some microscopy modalities such as HILO, which is limited by the distance of cell nuclei to the microscope objective. Given the advantages provided by the two-dimensional iPSC/iTSC co-culture model for live single molecule tracking microscopy described here, this model could offer a way to study the dynamics of additional transcription factors and signalling proteins that drive cell fate choices in early embryos.

Certain cancers are driven by oncogenic fusions with Hippo pathway transcription regulators, i.e. YAP, TAZ or TEADs (*32*). Interestingly, our high-resolution imaging studies showed that one such cancer fusion, YAP-TFE3, is less mobile and binds DNA longer than either YAP or TEAD1 and, like YAP, its nuclear behaviour is highly dependent on TEADs. Single molecule tracking revealed that TFE3 confers some DNA binding ability to YAP-TFE3, which is consistent with published genomic studies (*50*). YAP-TFE3 also slowed TEAD1’s nuclear mobility, indicating that it may promote TEAD DNA target search and participate in cooperative DNA binding, for example, by the DNA binding domains of both TEADs and TFE3. This suggests that other transcription factors that form fusion proteins that underpin different cancers also possess nuclear behaviour and DNA binding properties that differ from their endogenous proteins and that they will influence the DNA association kinetics of their interaction partners, although this remains to be tested.

During the course of our studies, we noted that the chromatin binding times of both YAP and TEAD1 occurred along a broad continuum, with a predominance of short binding events occurring for less than a second, and rarer long binding events occurring for up to a minute or more.

Therefore, we examined whether this could be explained by YAP and TEAD1 binding chromatin on different timescales in discrete nuclear sub-domains, paying close attention to nuclear condensates, given both YAP and TAZ form condensates that have been linked to transcription activation (*26–29*). Indeed, we found that TEAD1 was less mobile, and its dwell times were substantially higher in condensates than other regions of the nucleus, again consistent with the notion that extended YAP/TEAD chromatin association correlates with active transcription. Interestingly, by acutely disrupting the YAP/TEAD protein-protein interaction using a small molecule (IAG933) (*49*), we found that YAP was completely dependent on TEAD binding to localise to nuclear condensates. By contrast, acute IAG933 treatment actually increased the size and number of TEAD1 condensates and caused TEAD1 DNA dwell times to increase in nuclear condensates. While at first glance, this seems at odds with our observation that longer YAP/TEAD chromatin dwell times are associated with transcription activation, it is important to note that TEAD is a default repressor of transcription (*10, 11*), and that chromatin binding by TEADs and the VGLL4 transcription co-repressor is favoured upon IAG933 treatment (*49*). Thus, we postulate that the extended TEAD1 DNA binding times observed in nuclear condensates upon IAG933 treatment reflect the transcription repression function of TEAD1/VGLL4. For example, TEAD1 might bind longer to chromatin upon acute IAG933 treatment because the TEAD1/VGLL4 complex recruits chromatin regulatory proteins to establish a repressive chromatin environment. In future studies, it will be interesting to determine whether this is a general feature of transcription factors that, like TEADs, possess dual activation/repression roles depending on their cofactor binding.

Another question raised by our and other studies that have revealed that transcription factor-DNA binding coincides with transcription activation is how exactly does a transcription co-factor influence DNA binding time of its cognate transcription factor? Several possibilities exist; for example, a transcription co-factor could allosterically increase the affinity of transcription factors for their cognate DNA binding motifs. Alternatively, a recent study showed that nascent mRNA can form a physical complex with transcription factors and stabilise their interactions with DNA (*54*). Additionally, this phenomenon could be explained by proteins that can form phase separated condensates through intrinsically disordered regions (*55*). As detailed, here and previously (*27*), such proteins exhibit reduced mobility inside condensates, which may promote DNA target search and could theoretically extend the chromatin binding time of their partner proteins.

### Limitations

The main microscopy approach used in this study - single molecule tracking - is limited by photobleaching, even with the use of photostable fluorescent dyes and gentle imaging modalities like HILO (*35*). Bleaching will have a disproportionate impact on long-lived DNA binding events; one way to alleviate this is with the use of photobleaching correction methods (*37*), as employed here, but the longest DNA binding events are likely to still be underestimated. Further, whilst we studied protein mobility in scenarios that are known to robustly modulate Hippo-regulated transcription, we did not assess transcription of individual gene loci in real time and binding of individual YAP and TEAD1 molecules specifically to them. Finally, given the very high degree of homology and similar behaviour in numerous functional studies (*56*), we chose to study only one TEAD family protein (i.e. TEAD1, but not TEAD2, 3 or 4). It is unlikely, but formally possible that TEAD2, 3 and/or 4 exhibit different biophysical behaviours than TEAD1.

## MATERIALS AND METHODS

### Plasmids

All HaloTag plasmids (both N- and C-terminally tagged) were created by cloning cDNAs into either a pReceiver-M49-HaloTag plasmid (for transient transfections) or a plenti-DEST-CMV-Neo plasmid for Doxycycline-inducible stably expressed plasmids. All GFP plasmids were created by cloning cDNAs into pEGFP-N1. The HA-TEAD1 plasmid created by cloning the TEAD1 cDNA into plenti-DEST-CMV-Neo . All mutations were engineered using PCR-based mutagenesis strategies.

### Cell culture

HeLa and HEK293T cells were cultured in DMEM (Thermo Fisher Scientific) supplemented with 10% Fetal Bovine Serum (Hyclone, SH30396.03). MCF10A cells were cultured in DMEM + F12 supplemented with Horse Serum (Life technology), Epidermal Growth Factor Human (Sigma-Aldrich, SRP3027-500UG), Hydrocortisone (Sigma-Aldrich: H0135-1MG), Cholera Toxin (Sigma-Aldrich, C8052-2MG) and Insulin (Actrapid Penfill, 100 units/ml). iPS cells were reprogrammed and maintained in a feeder-free E8 media (Thermo Fisher, A1517001) with vitronectin (VTN-N, Thermo Fisher, A14700) as published (*57*). Cells were cultured in 37 °C, 5% O_2_ and 5% CO_2_, and media was changed daily. Cells were passaged every 3-5 days with 0.5mM EDTA (Thermo Fisher, 15575-038). iTSC’s were reprogrammed and maintained in collagen IV (Sigma-Aldrich, C5533) coated plates in the following medium, as published (*57*). DMEM/F-12 GlutaMAX (Thermo Fisher, 10565018) with 0.3% (wt/vol) BSA (Sigma-Aldrich, A9576), 0.2% (vol/vol) FBS (HyClone, SH30071.03), 1% (vol/vol) ITS-X supplement (Thermo Fisher, 51500056), 0.1 mM 2- mercaptoethanol (Thermo Fisher, 21985023), 0.5% (vol/vol) pen/strep Thermo Fisher, 15140122), 1.5 μg/ml L-ascorbic acid (Sigma-Aldrich, A4544), 5 μM Y27632 (Selleckchem, S1049), 2 μM CHIR99021 (Sigma-Aldrich, SML1046), 0.5 μM A83-01 (Sigma-Aldrich, SML0788), 1 μM SB431542(Selleckchem, S1067), 50 ng/ml EGF (Peprotech, AF-100-15) and 0.8 mM VPA (Sigma-Aldrich, P4543). Cells were cultured in 37 °C and 5% CO_2_ and media were changed every other day. Cells were passaged every 4-5 days with Typle express (Thermo Fisher, 12604021)

### Viral transduction of cell lines

HEK293T cells were transfected using either polyethylene (PEI) transfection or LTX methods to generate lentiviral particles to transduce Halo-YAP/TEAD1 expression vectors. A lentiviral packaging mixture was prepared by combining OptiMEM, psPAX2, pMD2.G, and the lentiviral expression vector. iPSC and iTSC were plated at a density of 5 × 10^4^ cells per well in a 12-well plate, after incubation at 37 °C for 24 h, transduction was performed with a multiplicity of infection (MOI; the ratio of infectious virions to cells) of 1 and 5, respectively, at a final volume of 0.5 ml per well with culture medium in the presence of polybrene transfection reagent (1:1700; TR-1003-G; EMD Millipore). The culture medium was replaced with 1 mL of culture medium every 24 h. After 72 h, transduced cells were passaged or sorted on the BD influx system flow cytometer. GFP positive cells from the cultured iPSC Halo-YAP, iPSc Halo-TEAD1, iTSC Halo-YAP and iTSC Halo-TEAD1 were collected and cultured.

### iPS/iTS cell co-culture

For Halo-YAP or Halo-TEAD1, iTS cells were seeded at a density of 30 × 10^4^ cells per dish in the imaging dishes (ibidi, 81156) coated with Collagen IV. After 48h, media were removed and 40 × 10^4^ of iPS cells were added in E8 medium with Y-27632 (10 µM), respectively. Doxycycline (2 µg/ml, Sigma 33429-100MG-R) was also added, and cells were cultured in 37 °C and 5% CO2 for 24h prior to imaging. Phenol red free DMEM/F12 (Thermo Fisher 11039-021) supplemented with E8 supplement was used during image.

### Immunoblotting

Whole-cell lysates were prepared in RIPA buffer and subjected to SDS PAGE electrophoresis. Proteins were transferred to PVDF membranes (Millipore) and probed with primary antibodies specific for the following proteins: YAP (Cell Signaling 4912), Phospho-YAP (Ser127; Cell Signaling 4911), TEAD1 (BD Biosciences 610922), or Tubulin (T9026 Sigma-Aldrich), followed by detection with secondary antibodies and chemiluminescence visualization.

### Immunofluorescence

iTS and iPS cells fixed in 4% paraformaldehyde were immunostained with mouse anti-YAP monoclonal primary antibodies (Abnova, H00010413-M01) at 1:100 dilution. Secondary antibodies conjugated to either Alexa Fluor 568 or Alexa Fluor 647 (Invitrogen) were used at a concentration of 1:500. Hoechst 33342 (1 ug/ml) was used to stain nuclei.

### Single Molecule Tracking Microscopy

To obtain optimal single molecule staining of Halo-YAP or Halo-TEAD1, cells were incubated with ∼2-5 nM of Halo JF549 ligand (Promega) in their media for ∼15-30 mins prior to imaging. In experiments to label all Halo-tagged molecules, Halo JF549 or Halo JFX650 ligand were applied at a concentration of 100 nM (e.g. Fig. 6 and fig. S8). In most experiments, DNA was simultaneously stained using Hoechst 33342 (NucBlue™ Live ReadyProbes™ Reagent, Invitrogen). Cells were maintained at 37°C with 5% CO_2_ on the microscope using a Tokai Hit incubation system. SMT was carried out on a Zeiss Elyra microscope using a Zeiss α Plan-Apochromat (100×/1.46 NA OIL DIC VIS) objective. The HILO imaging modality was used in combination with a high power TIRF field (TIRF_HP) and an additional 1.6× OptoVar Lens. TIRF angles of 57° (iPS/iTS cells), 58° (MCF10A cells), or 59° (HeLa cells) were used to optimally section the nuclear plane with HILO. Halo JF549 ligand was excited using a 200 mW 561 nm laser using either 10% (Fast SMT) or 5% (Slow SMT) laser power with emission light passed through 570-620 nm bandpass filter and detected using an Andor iXon 897 EMCCD camera. Hoechst 33342 was visualised using a 50 mW 405 nm laser with emission light passed through a 420-480 nm bandpass filter to the camera. When bulk Halo-tagged TEAD1 was imaged, Halo-JFX650 was excited using a 150 mW 642nm laser and emission light passed through a 650 nm longpass filter to the camera. The microscopy parameters, lasers, and cameras were controlled through Zen 2012 SP5 (Black, Version 14.0.0.0). Using these parameters, we performed two different acquisition techniques; Fast SMT, which uses a 20 ms acquisition speed to acquire 6000 frames without intervals, and Slow SMT, which uses a 500 ms acquisition speed to acquire 500 frames without intervals. The only exception to this was our analysis of TEAD1 condensates (Fig. 6. And fig. S8). In this case, 3000 frames were acquired for Fast SMT, and 250 frames for Slow SMT, before and after IAG933 treatment, in the same cells.

### Fast Single Molecule Tracking

Analysis was performed as in (*58*). Briefly, raw Fast (20 ms) SMT data was analysed using the PalmTracer plugin for Metamorph (*36, 59*). PalmTracer was used to localize and track molecules in order to obtain their trajectories, and to calculate the mean square displacement (MSD), and diffusion coefficient (D) for each trajectory. For molecule localisation, we used a watershed of size 6. To reduce non-specific background, and to reduce the likelihood of mistracking, trajectories were filtered based on a minimum length of 8 and a maximum length of 1000, and a maximum travel distance of 5 μm. For visualisation of trajectories, a zoom of 8 with a fixed intensity and size of 1 was used. A spatial calibration of 100 nm and a temporal calibration of 20 ms was used.

Analysis files produced by PalmTracer were then used as input for AutoAnalysis_SPT software (https://github.com/QBI-Software/AutoAnalysis_SPT/wiki). AutoAnalysis_SPT compiles the results obtained for each cell to obtain the average MSD, calculates the average area under the curve (AUC) of the MSD for each cell, generates a histogram showing the distribution of the different Log_10_ Diffusion Coefficients, and calculates the mobile to immobile ratio for each cell. Here, we used 10 MSD points, with a time interval of 0.02 s (20 ms acquisition time) and included trajectories with a minimum Log_10_ Diffusion Coefficient of −5, and a maximum Log_10_ Diffusion Coefficient of 1. For the Log_10_ Diffusion Coefficient histogram, a bin width of 0.1, and a mobile to immobile threshold of −1.45 (∼0.035 μm^2^/sec) was selected based on previous studies (*58*).

### Slow Single Molecule Tracking

Images were acquired with an exposure time of 500ms such that fast-moving molecules were blurred out and only immobile, DNA-bound molecules were observed. Single molecule localisation and tracking was performed using SLIMfast for MATLAB (Version 2015a), as in (*34, 58, 60, 61*). SLIMfast uses a modified version of the multiple-target tracing (MTT) algorithm (*62*). SLIMfast batch processing was performed using an error rate of 10^−7^, a detection box of 7 pixels, maximum number of iterations of 50, a termination tolerance of 10^−2^, a maximum position refinement of 1.5 pixels, an NA of 1.46, a PSF scaling factor of 1.35, and 20.2 counts per photon, an emission of 590 nm, a lag time of 500 ms and a pixel size of 100 nm. Trajectories were filtered with the maximum expected diffusion coefficient of 0.1 μm^2^/s. Dwell time data was compiled for each condition and a frequency distribution was generated using GraphPad Prism. Dwell time data was subsequently photobleach corrected and analysed in MATLAB (Version 2024a). To exclude the effect of the noisy tail of the survival distribution, timepoints with fewer than 30 cumulative events were excluded from fitting, except in the case of Fig. 6, where all values were included in the fitting. As described in (*37, 39*), the photobleaching rate of Halo JF549 ligand was determined using SMT of Halo-H2B, expressed in HeLa cells, and acquired using different TIRF angles to reflect different experimental settings. Survival distributions were fit to bi- or triple-exponential functions, and the best fit was determined using Bayesian Information Criterion (BIC). The smallest exponential component, corresponding to the bleaching rate of JF549 was used for subsequent correction of Slow tracking SMT data. Bleach-corrected survival distributions derived from tracking data of Halo-tagged proteins were fit to a bi-exponential fit or a power-law fit.

### Confocal Microscopy

Live confocal microscopy of Halo-TEAD1 and YAP-GFP (Fig. 6) was performed using a Zeiss Elyra LSM780 microscope, equipped with a Zeiss C-Apochromat (63×/1.2 NA W Korr UV-VIS-IR) objective, 488nm (GFP) or 561nm (Halo-JF549) lasers, and a spectral GaAsP detector with two flanking PMT’s (detection windows used were 490-553 nm for GFP and 562-641 nm for Halo- JF549). Cells were maintained at 37°C with 5% CO_2_ on the microscope using a Tokai Hit incubation system. For LATSi experiments (Fig. 4), cells were first treated with either DMSO or 3 μM LATSi (*45*) for 2 hrs prior to imaging. For IAG933 experiments (Fig. 6), a 10X solution of IAG933 (*49*) in imaging media was added to cells (to a working concentration of 3 μM) during microscopy and immediately imaged.

### Raster Image Correlation Spectroscopy

All RICS experiments were performed on an Olympus FV3000 laser scanning microscope. A 60× water immersion objective 1.2 NA was used for all experiments and live HeLa or MCF10A cells were imaged at 37°C in 5% CO_2_. For RICS experiments on Halo-YAP or Halo-TEAD1 (Fig. 1F and Fig. S5B), cells were stained for 15 mins with 1 μM of JF549 Halo-tag dye to label all molecules, where JF549 was excited by a solid-state laser diode operating at 561 nm. The JF549 emission was collected through a 550 nm long pass filter by an external photomultiplier detector (H7422P-40 of Hamamatsu) fitted with a 620/50 nm bandwidth filter. A 100-frame scan acquisition of the JF549 signal was collected by selecting a region of interest within a nucleus at zoom 20, with a 1 Airy unit pinhole size which for a 256 × 256-pixel frame size resulted in a pixel size of 41 nm. The pixel dwell time was set to 12.5 μs, which resulted in a line time of 4.313 ms and a frame time of 1.108 s. RICS analysis was carried out using SimFCS software (Globals), using a 10-frame moving average subtraction. This involved application of the RICS function to each time series acquisition and extraction of the apparent diffusion coefficient (D) by fitting each resulting 3D RICS profile to a 1-component diffusion model.

### Statistical Analyses

GraphPad Prism (Version 10.1.2) was used to generate graphs and to perform statistical analyses.

## ACKNOWLEDGEMENTS

We thank members of the Harvey lab for discussions and comments on the manuscript. We thank Xiaomeng Zhang for technical support, and Jean Baptiste Sibarita and Fred Meunier for making available the PalmTracer plugin for Metamorph and AutoAnalysis_SPT software. K.F.H was supported by a Senior Research Fellowship (APP1078220) and Investigator grant (APP1194467) from the National Health and Medical Research Council of Australia (NHMRC). E.H was supported by a Future Fellowship (FT200100401), Discovery Project (DP210102984), Centre of Excellence (CE230100021), and LIEF Project (LE210100046) from the Australian Research Council (ARC). This research was supported by the ARC (DP180102044, DP190101743, DP22010523 and DP230101406) and the NHMRC (APP1157737). We acknowledge the Peter Mac Centre for Advanced Histology and Microscopy and support to them from the Peter MacCallum Cancer Foundation and the Australian Cancer Research Foundation. We acknowledge the Monash Micro Imaging Facility.

