## Supplementary data for "Hippo signalling regulates the nuclear behaviour and DNA dwell times of YAP and TEAD to control transcription"

### SUPPLEMENTARY MATERIALS

### SUPPLEMENTARY FIGURES

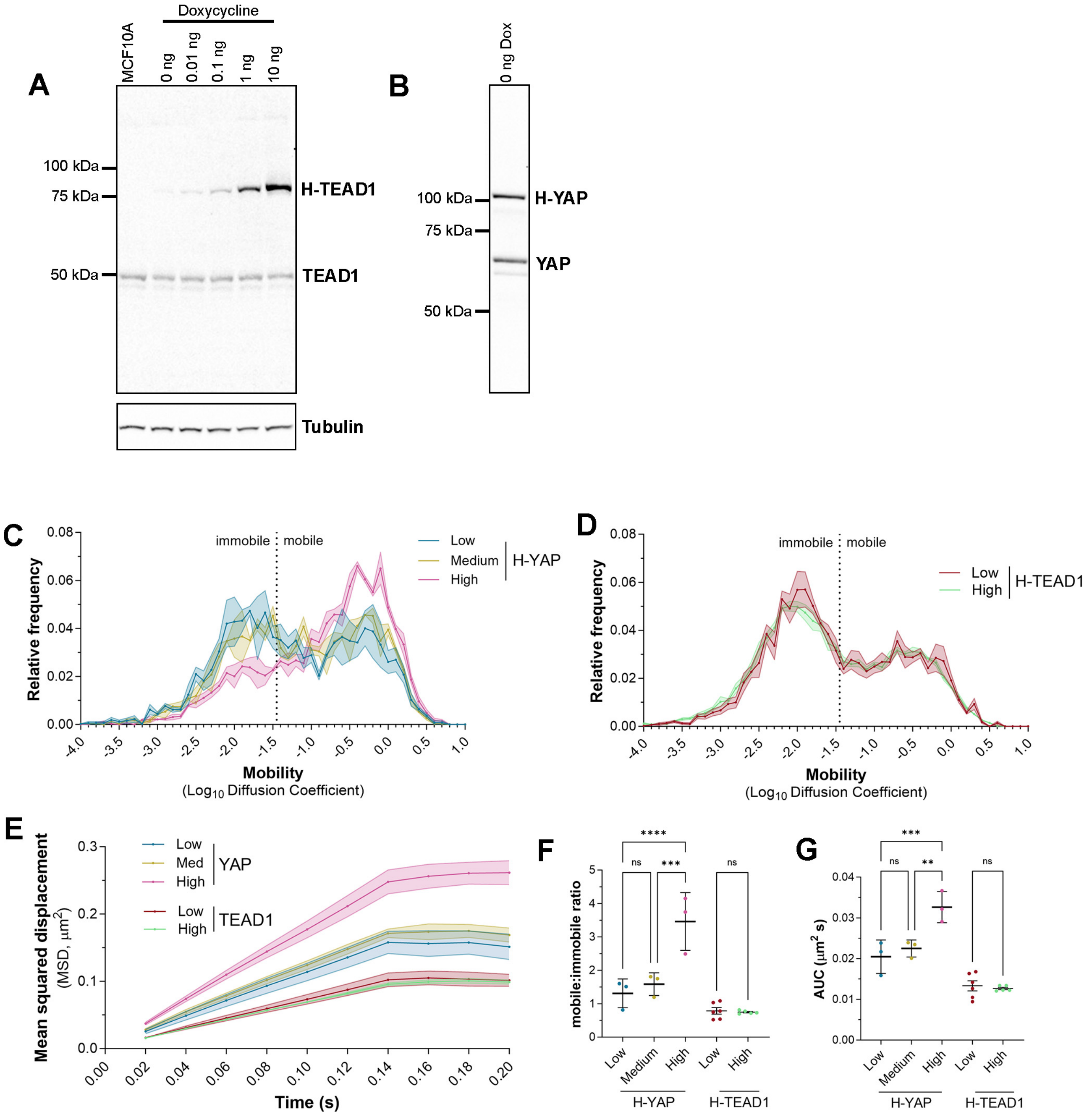


**Supplementary Figure 1. Analysis of the impact of expression level on YAP and TEAD1 biophysical properties in cell nuclei.**

**A and B)** Immunoblots of lysates from MCF10A cells stably transduced with vectors expressing either HaloTag-TEAD1 (A) or HaloTag-YAP (B). Cells were either untreated, or treated with the indicated concentrations of Doxycycline for 24h. Membranes were probed with anti-TEAD1 (A) or anti-YAP (B) and equal protein loading in (A) was confirmed by detection of Tubulin.

**C and D)** Charts showing the relative frequency of molecule mobilities ($Log_{10}$ diffusion coefficient) for YAP (C) and TEAD1 (D) molecules in MCF10A cell nuclei over time. Data presented as the mean ± SEM. Mobile and immobile fractions are indicated. Low (blue), medium (olive) and high (pink) expression levels were assessed for YAP, and low (red) and high (green) for TEAD1.

**E)** Chart showing the mean squared displacement ($\mu m^2$) of nuclear YAP and TEAD1 molecules in MCF10A cell nuclei expressing different levels of YAP or TEAD1 protein over time. Data presented as the mean ± SEM.

**F and G)** Charts showing the mobile to immobile ratio (F) and the area under the curve (G, $\mu m^2/s$) of TEAD1 and YAP molecules in MCF10A cell nuclei. Data presented as mean ± SD, **** $p < 0.0001$, *** $p < 0.001$, ** $p < 0.01$ ns: not significant, n = 3 for YAP at each expression level, and n = 6 and 5 for low and high levels of TEAD1, respectively.

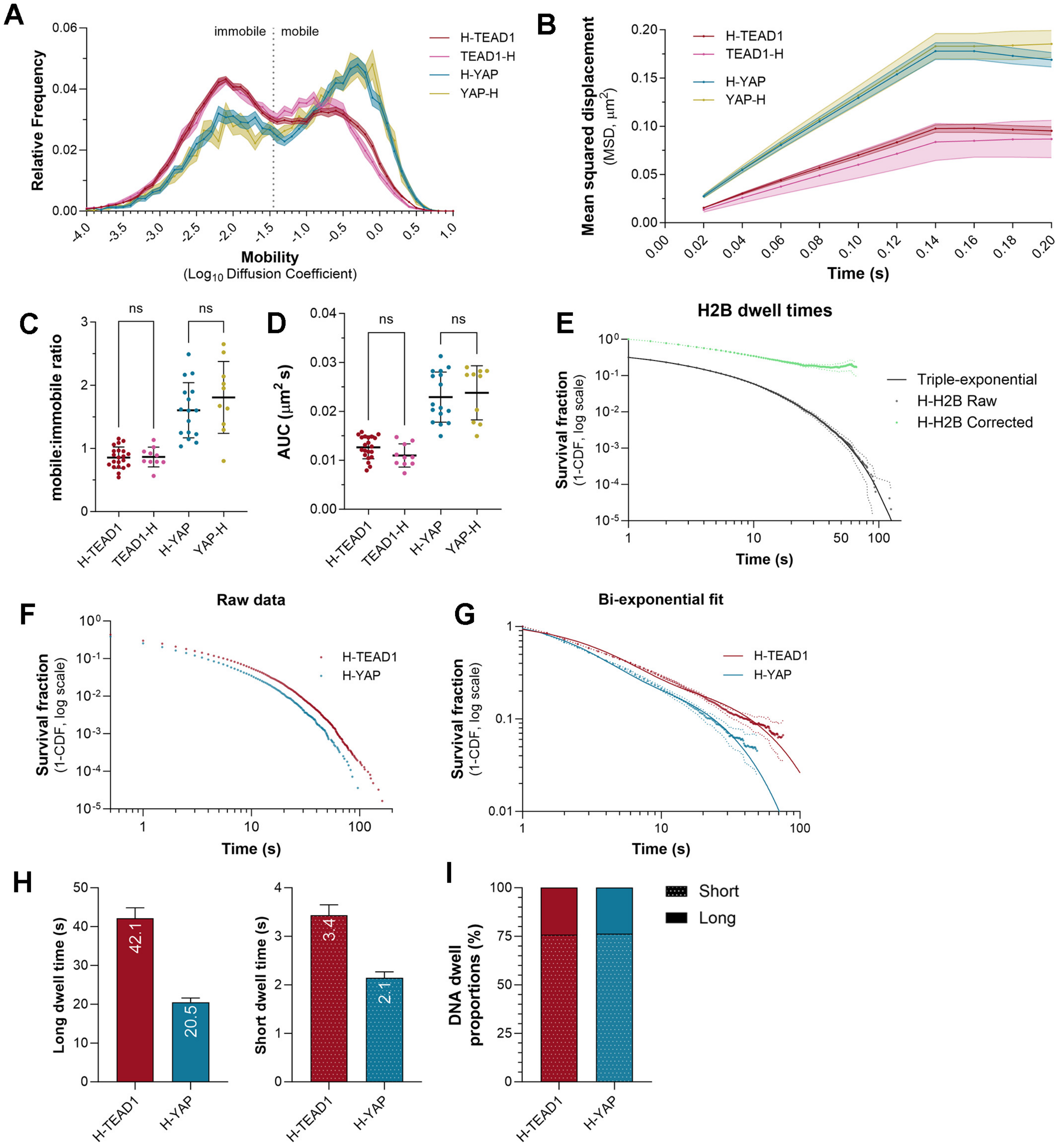


**Supplementary Figure 2. Analysis of the impact of the position of the HaloTag YAP and TEAD1 biophysical properties in cell nuclei.**

**A)** Chart showing the relative frequency of molecule mobilities ($Log_{10}$ diffusion coefficient) for YAP (C) and TEAD1 (D) molecules in MCF10A cell nuclei over time. Both YAP and TEAD1 were tagged at either the N- or C-termini with HaloTag (H). Data presented as the mean ± SEM. Mobile and immobile fractions are indicated.

**B)** Chart showing the mean squared displacement ($\mu m^2$) of N- and C- terminus tagged YAP or TEAD1 molecules in MCF10A cell nuclei over time. Data presented as the mean ± SEM.

**C and D)** Charts showing the mobile to immobile ratio (C) or the area under the curve (D, $\mu m^2/s$) of TEAD1 and YAP molecules in MCF10A cell nuclei. Both YAP and TEAD1 were tagged at either the N- or C-termini with HaloTag. Data presented as mean ± SD, p-values obtained using an unpaired t-test; ns: not significant, n = 16 and 10 cells for H-YAP and YAP-H, and n = 21 and 10 cells for H-TEAD1 and TEAD1-H.
**E)** Chart showing the survival distribution of Halo-tagged Histone H2B molecules in HeLa cell nuclei. Raw data and photobleach-corrected data are shown ± 99% C.I. (dotted lines). Solid line is the triple-exponential fit to the raw data. n = 47,997 trajectories from 17 cells.
**F)** Chart showing the raw survival distribution of YAP (blue) and TEAD1 (red) molecules in MCF10A cell nuclei. n = 61,860 trajectories from 31 cells for TEAD1, and 27,748 trajectories from 20 cells for YAP.
**G)** Chart showing the photobleach-corrected survival distribution of Halo-TEAD1 or Halo-YAP molecules in the nuclei of MCF10A cells, with bi-exponential fitting (solid lines). Dotted lines are ± 99% CI.
**H and I)** Charts showing long and short DNA dwell times (H) and relative proportion of long and short DNA dwell times (I) for Halo-tagged TEAD1 and YAP. Dwell times are calculated as 1/k from the bi-exponential fit and displayed ± 95% CI.

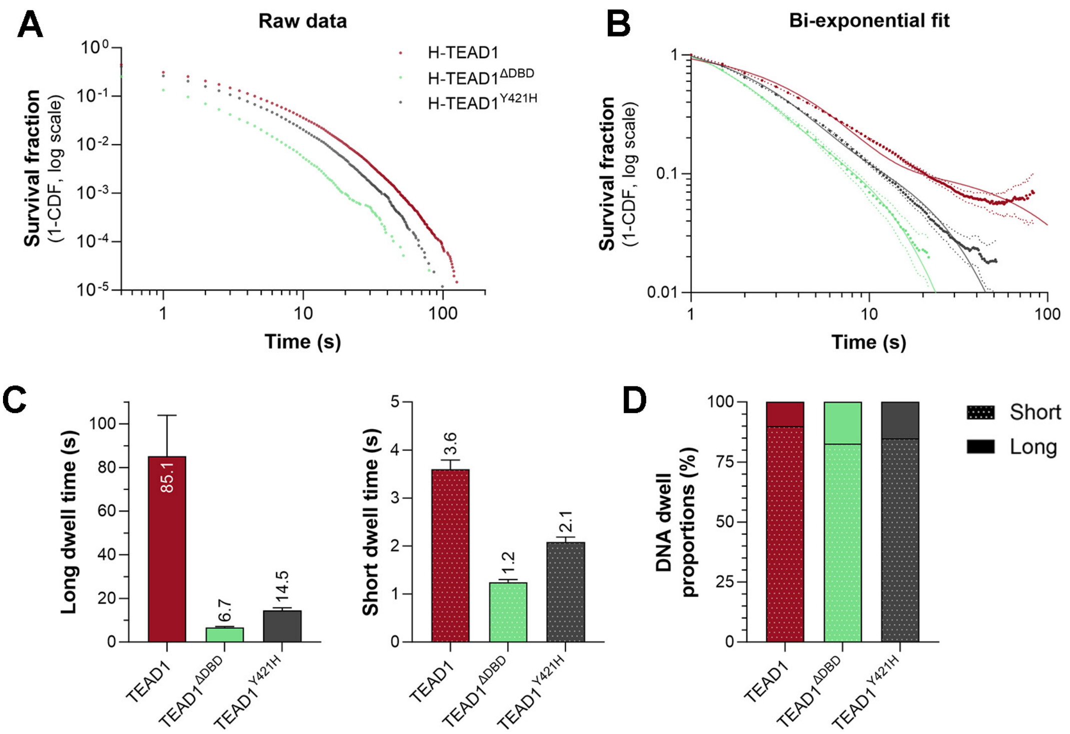


**Supplementary Figure 3. The impact of different mutations on TEAD1 nuclear behaviour.**

**A)** Chart showing the raw survival distribution of wild-type and mutant TEAD1 proteins in HeLa cell nuclei. n = 202,613 trajectories from 26 cells for TEAD1, 38,766 trajectories from 22 cells for TEAD1$^{\Delta DBD}$, and 83,778 trajectories from 21 cells for TEAD1$^{Y421H}$.

**B)** Chart showing the photobleach-corrected survival distribution of wild-type and mutant TEAD1 molecules in the nuclei of HeLa cells, with bi-exponential fitting (solid lines). Dotted lines are ± 99% CI.

**C and D)** Charts showing the mean long and short DNA dwell times (C) and relative proportion of long and short DNA dwell times (D) for TEAD1, TEAD1$^{\Delta DBD}$, and TEAD1$^{Y421H}$ molecules, as derived from the bi-exponential function. Dwell times are calculated as 1/k from the bi-exponential fit and displayed ± 95% CI.

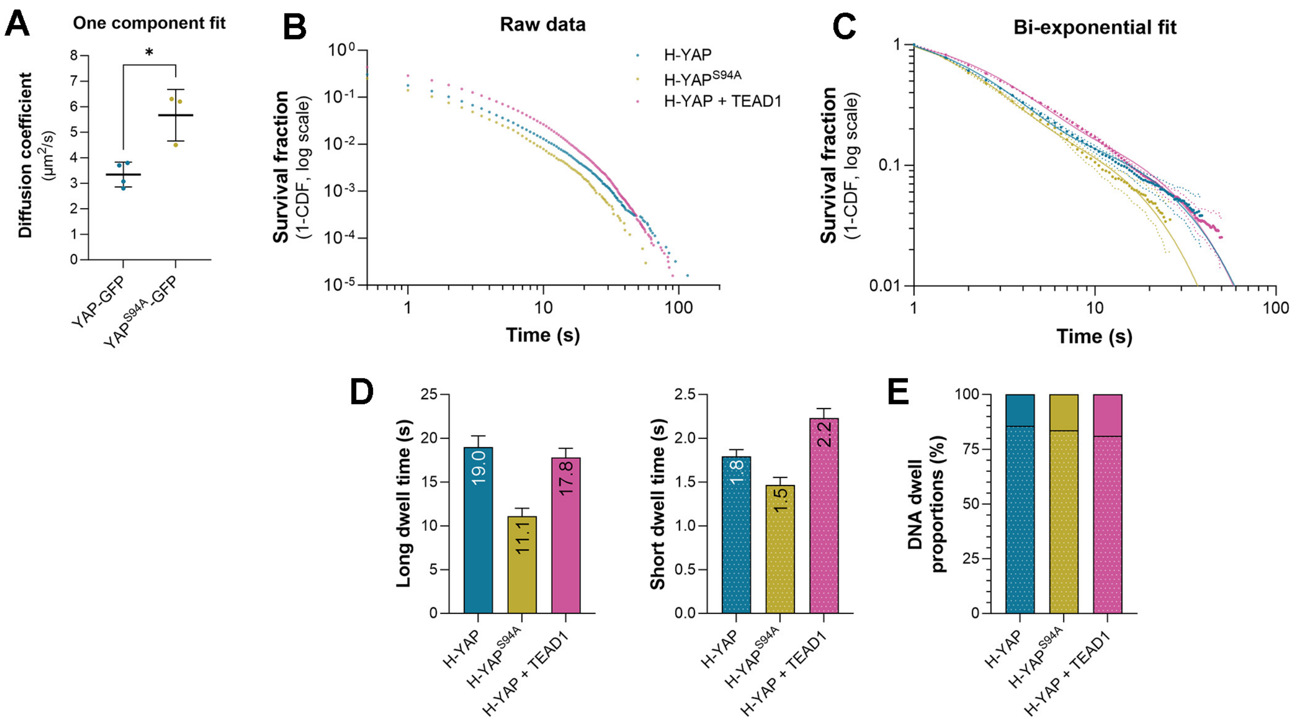


**Supplementary Figure 4. TEAD1 profoundly influences YAP nuclear behaviour.**

**A)** Chart of diffusion coefficient (μm$^2$/s) of GFP-tagged YAP and YAP$^{S94A}$ in HeLa cell nuclei, as determined by fluorescence correlation spectroscopy. Data is fit to a one-component model, and presented as mean ± SD, * $p < 0.05$, n = 4 and 3 cells for YAP-GFP and YAP$^{S94A}$-GFP, respectively.

**B)** Chart showing the raw survival distribution of YAP (blue), YAP$^{S94A}$ (olive), and YAP in the presence of TEAD1 overexpression (green), in HeLa cell nuclei. n = 62,126 trajectories from 24 cells for YAP, 33,571 trajectories from 13 cells for YAP$^{S94A}$, and 126,127 trajectories from 16 cells for YAP with overexpressed TEAD1.

**C)** Chart showing the photobleach-corrected survival distribution of wild-type and mutant YAP molecules, and YAP in the presence of TEAD1 overexpression, in HeLa cell nuclei. Bi-exponential fits are shown (solid lines). Dotted lines are ± 99% CI.

**D and E)** Charts showing the mean long and short DNA dwell times (D) and relative proportion of long and short DNA dwell times (E) for wild-type and mutant YAP proteins and YAP in the presence of TEAD1 overexpression, as derived from the bi-exponential function. Dwell times are calculated as 1/k from the bi-exponential fit and displayed ± 95% CI.

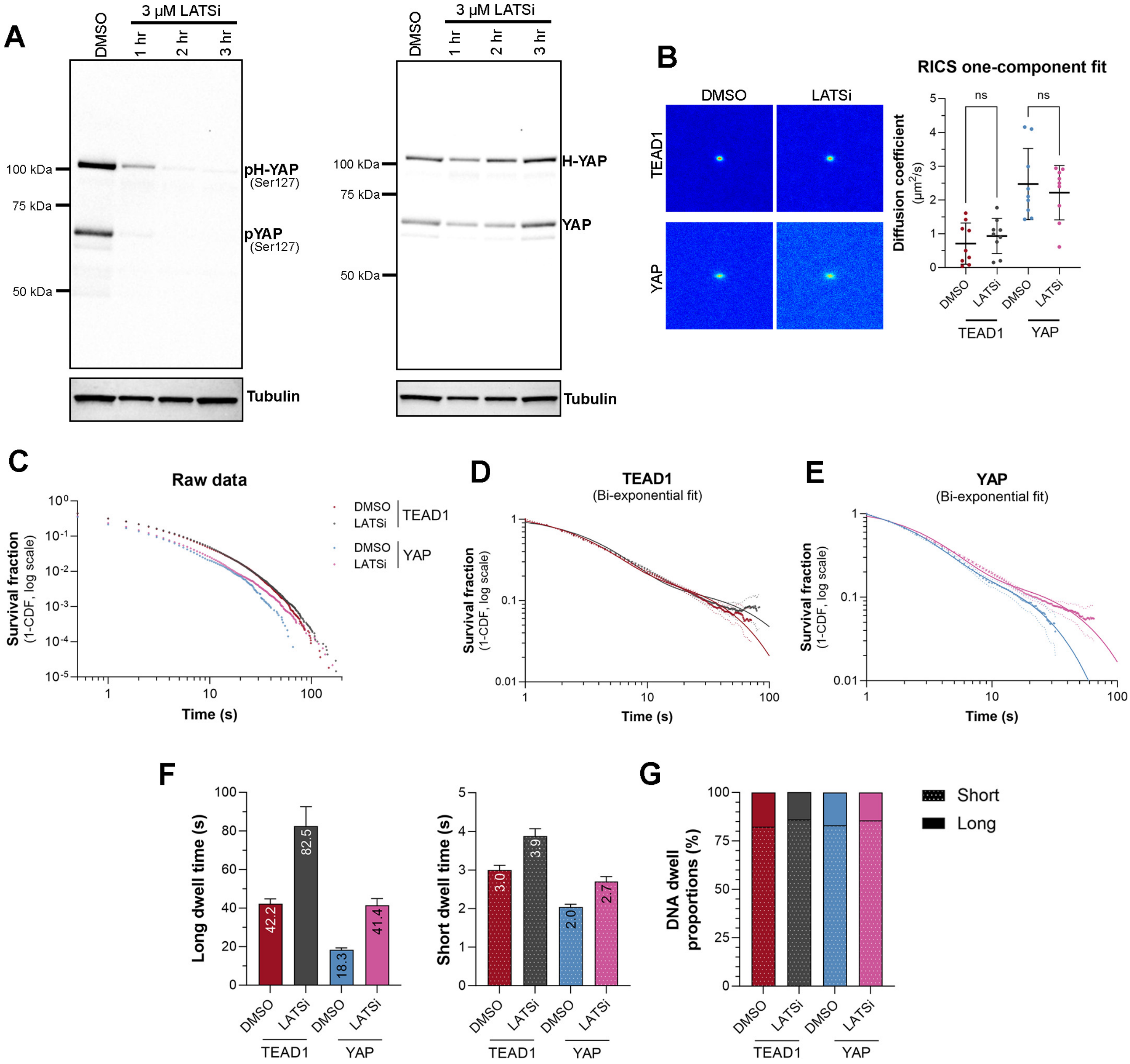


**Supplementary Figure 5. Hippo signalling limits the DNA dwell times of YAP and TEAD.**

**A)** Immunoblots of lysates from MCF10A cells stably transduced with HaloTag-YAP (B). Cells were either untreated, or treated with 3 μM LATS inhibitor for the indicated durations. Membranes were probed with anti-phospho-S127 YAP (left blot) or anti-YAP (right blot) and equal protein loading was confirmed by detection of Tubulin.

**B)** On the left are three-dimensional RICS correlation functions derived from MCF10A cells expressing Halo-tagged YAP or TEAD1 with or without LATSi treatment. On the right is a chart of the diffusion coefficients (μm$^2$/s) of YAP and TEAD1 with or without LATSi treatment, as determined by fluorescence correlation spectroscopy. Data is fit to a single-component model, and presented as mean ± SD, ns: not significant, n = 9 cells for all conditions.

**C)** Chart showing the raw survival distribution of TEAD1 and YAP in MCF10A cell nuclei with or without LATSi treatment. n = 54,886 trajectories from 21 cells for TEAD1 (DMSO), 68,992 trajectories from 17 cells for TEAD1 (LATSi), 14,104 trajectories from 19 cells for YAP (DMSO), and 46,056 trajectories from 22 cells for YAP (LATSi).

**D and E)** Charts showing the photobleach-corrected survival distribution of TEAD1 (C) or YAP (D) molecules in MCF10A cell nuclei with or without LATSi. Bi-exponential fits are shown (solid lines). Dotted lines are ± 99% CI.

**F and G)** Charts showing the mean long and short DNA dwell times (F) and relative proportion of long and short DNA dwell times (G) of TEAD1 or YAP with or without LATSi treatment, as derived from the bi-exponential function. Dwell times are calculated as 1/k from the bi-exponential fit and displayed ± 95% CI.

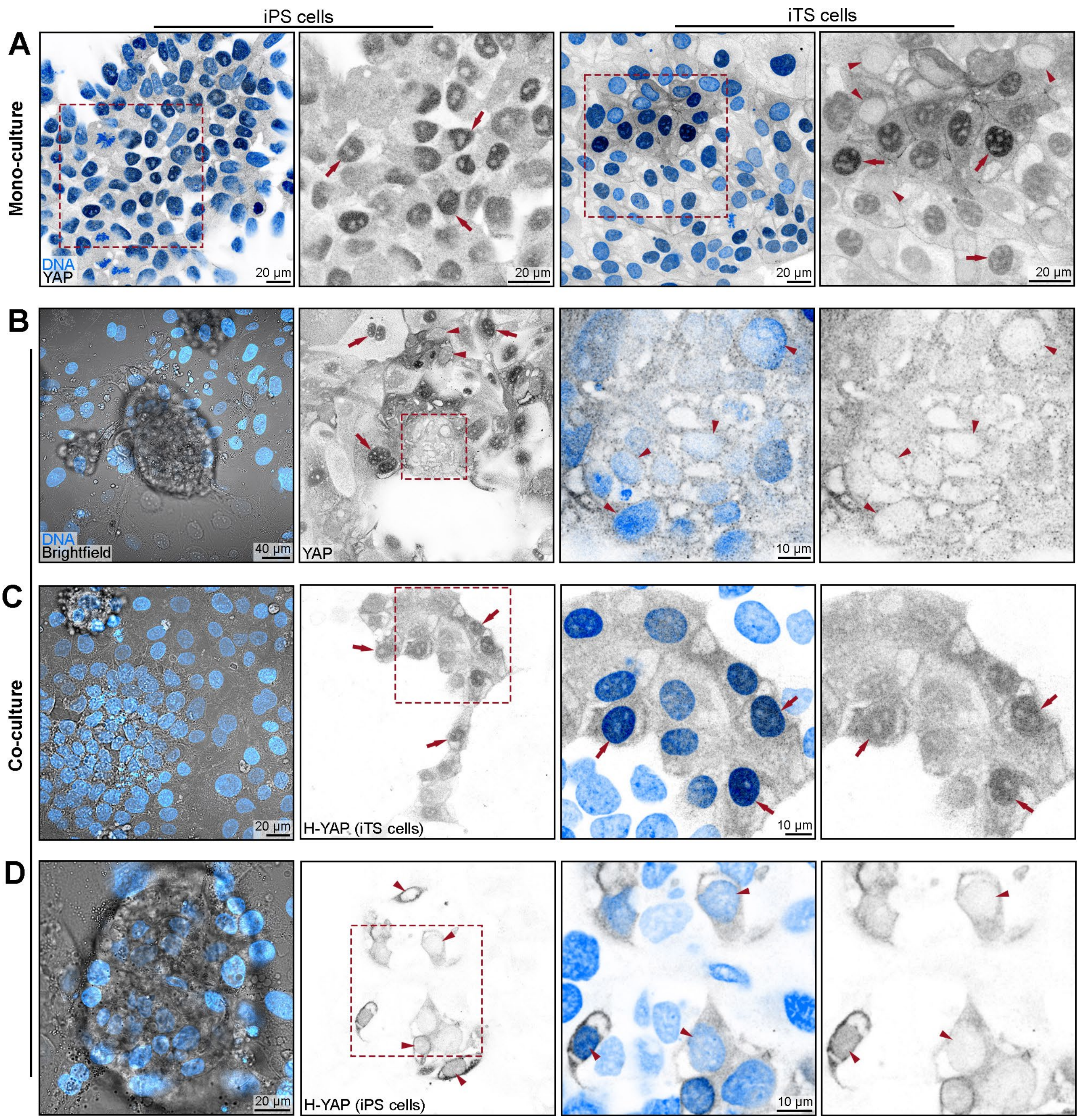


**Supplementary Figure 6. YAP subcellular localisation in different iPS/iTS cell culture models.**

**A)** Confocal microscope images of iPS cells and iTS cells grown as mono-cultures. DNA is labelled with Hoechst (blue), and antibody-detected YAP is grey. YAP was predominantly nuclear in both cell types; examples are indicated by red arrows. Some iTS cells had low nuclear YAP (red arrowheads). Red boxed regions are shown at higher magnification in the right panels, scale bars are indicated.

**B)** Confocal microscopy images of co-cultured wild-type iPS and iTS cells. DNA is labelled with Hoechst (blue), and antibody-detected YAP is grey. Brightfield shows a central sphere of iPS cells surrounded by a lawn of iTS cells. YAP is predominantly nuclear in many iTS cells (red arrows), and most iPS cells had low nuclear YAP (arrowheads). Red boxed region is shown at higher magnification in the right-hand panels. Scale bars are indicated.

**C)** Confocal microscope images of iTS cells expressing Halo-tagged YAP, co-cultured with wild-type iPS cells. DNA is labelled with Hoechst (blue), and Halo-YAP is detected by Halo-JF549 dye (grey). YAP is predominately nuclear in iTSCs, as indicated by red arrows. Red boxed region is shown at higher magnification in the right-hand panels. Scale bars are indicated.
**D)** Confocal microscope images of iPS cells expressing Halo-tagged YAP, co-cultured with wild-type iTS cells. DNA is labelled with Hoechst (blue), and Halo-YAP is detected by Halo-JF549 dye (grey). YAP is predominately cytoplasmic in iTSCs, as indicated by red arrows. Red boxed region is shown at higher magnification in the right-hand panels. Scale bars are indicated.

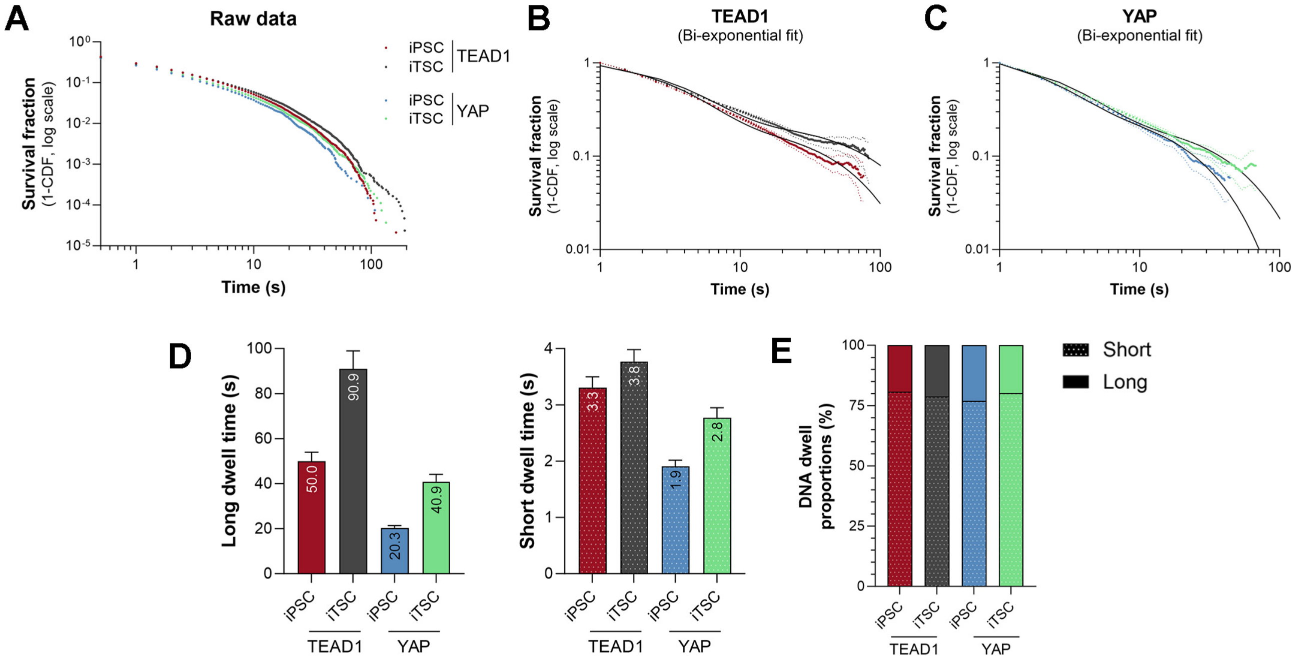


**Supplementary Figure 7. Longer YAP and TEAD DNA binding times in cells with intrinsically high YAP/TEAD activity.**

**A)** Chart showing the raw survival distribution of Halo-tagged TEAD1 and YAP molecules in iPSC or iTSC nuclei. n = 47,191 trajectories from 22 cells for TEAD1 (iPS cells), 42,339 trajectories from 22 cells for TEAD1 (iTS cells), 13,381 trajectories from 25 cells for YAP (iPS cells), and 26,931 trajectories from 23 cells for YAP (iTS cells).

**B and C)** Charts showing the photobleach-corrected survival distribution of TEAD1 (B) or YAP (C) molecules in iPSC or iTSC nuclei. Bi-exponential fits are shown (solid lines). Dotted lines are ± 99% CI.

**D and E)** Charts showing the mean long and short DNA dwell times (D) and relative proportion of long and short DNA dwell times (F) of TEAD1 or YAP in iPSC or iTSC nuclei, as derived from the bi-exponential function. Dwell times are calculated as 1/k from the bi-exponential fit and displayed ± 95% CI.

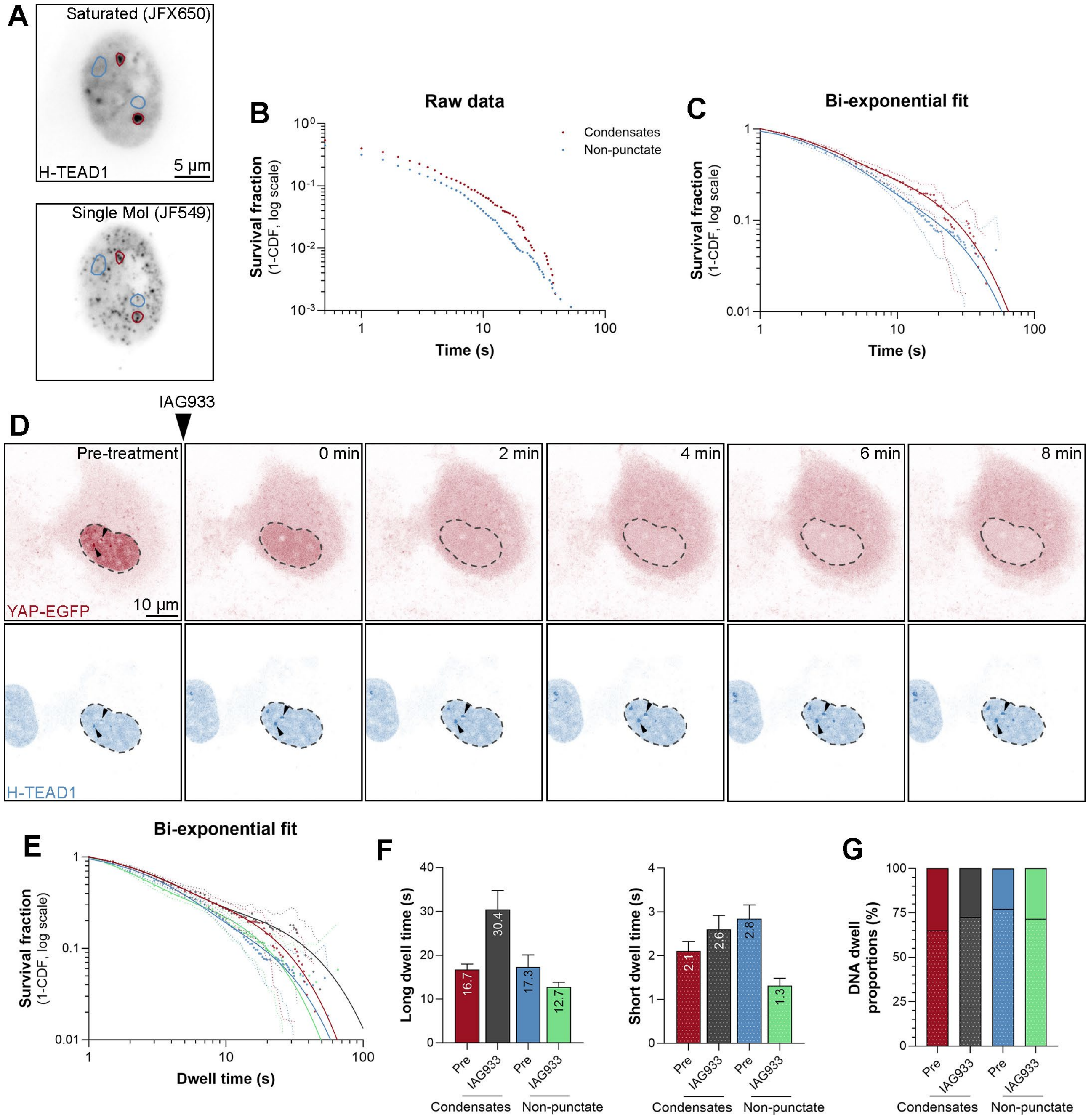


**Supplementary Figure 8. TEAD exhibits extended DNA binding behaviour in nuclear condensates**

**A)** Images of a single MCF10A cell nucleus expressing Halo-tagged TEAD1 (grey); saturating amounts of JFX650 labelled all Halo-TEAD1 protein (top panel), while a low concentration of JF549 was used to enable single molecule detection (bottom panel). Condensates (red ROI's) and non-punctate nuclear regions (blue ROI's) were selected based on images of the top panel and applied to single molecule tracking data collected from the bottom panel.

**B)** Chart showing the raw survival distribution of Halo-tagged TEAD1 in condensates and non-punctate nuclear regions. n = 1,080 trajectories in condensates, and 2,628 trajectories in non-punctate regions, from 14 cells.

**C)** Charts showing the photobleach-corrected survival distribution of TEAD1 in condensates and non-punctate nuclear regions of MCF10A cells, with bi-exponential fitting (solid lines). Dotted lines are ± 99% CI.

**D)** Images of a single HeLa cell expressing YAP-EGFP (red) and Halo-tagged TEAD1 (blue) pre- and post-treatment with 3 μM IAG933 every 2 minutes for 8 minutes. Arrows indicate TEAD1 condensates. Scale bars are indicated.

**E)** Charts showing the photobleach-corrected survival distribution of TEAD1 in condensates and non-punctate nuclear regions of MCF10A cells, before and after treatment with IAG933. Bi-exponential fits are shown (solid lines). Dotted lines are ± 99% CI.

**F and G)** Charts showing the mean long and short DNA dwell times (F) and relative proportion of long and short DNA dwell times (G) of TEAD1 in condensates and non-punctate nuclear regions of MCF10A cells before and after treatment with IAG933, as derived from the bi-exponential function. Dwell times are calculated as 1/k from the bi-exponential fit and displayed ± 95% CI.

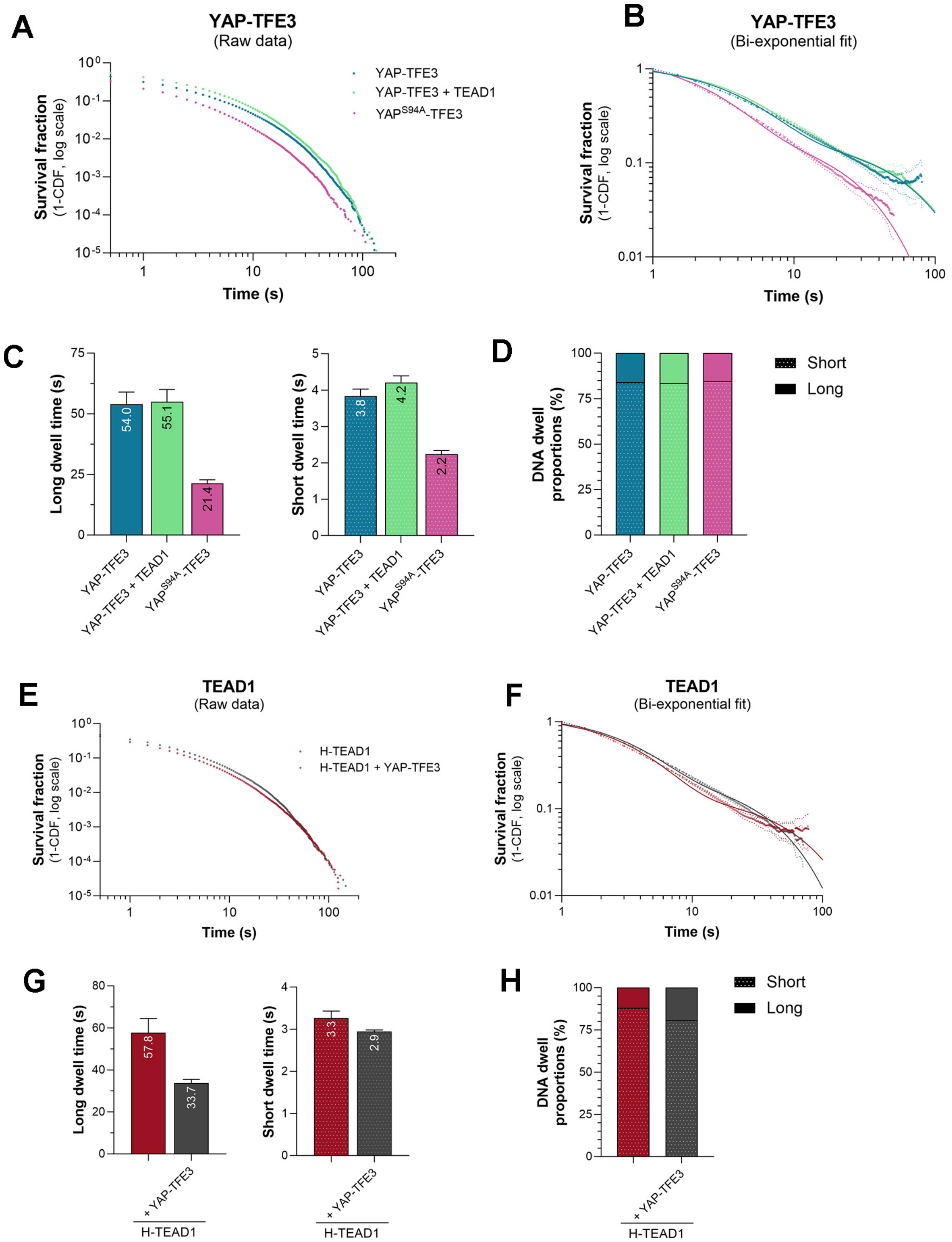


**Supplementary Figure 9. The nuclear behaviour of the YAP-TFE3 oncogenic fusion protein differs substantially from YAP but is still influenced by TEADs.**

**A)** Chart showing the raw survival distribution of YAP-TFE3 (blue), $YAP^{S94A}$-TFE3 (pink), and YAP-TFE3 in the presence of TEAD1 overexpression (green), in HeLa cell nuclei. n = 167,283 trajectories from 27 cells for YAP-TFE3, 104,009 trajectories from 19 cells for $YAP^{S94A}$-TFE3, and 86,983 trajectories from 20 cells for YAP-TFE3 with overexpressed TEAD1.

**B)** Chart showing the photobleach-corrected survival distribution of YAP-TFE3, YAP$^{S94A}$-TFE3, and YAP-TFE3 in the presence of TEAD1 overexpression, in HeLa cell nuclei. Bi-exponential fits are shown (solid lines). Dotted lines are ± 99% CI.
**C and D)** Charts showing the mean long and short DNA dwell times (C) and relative proportion of long and short DNA dwell times (D) for YAP-TFE3, YAP$^{S94A}$-TFE3, and YAP-TFE3 in the presence of TEAD1 overexpression, as derived from the bi-exponential function. Dwell times are calculated as 1/k from the bi-exponential fit and displayed ± 95% CI.
**E)** Chart showing the raw survival distribution of Halo-TEAD1 molecules, in the presence or absence of YAP-TFE3-GFP, in HeLa cell nuclei. n = 121,486 trajectories from 16 cells for TEAD1 alone, and 102,439 trajectories from 15 cells for TEAD1 with overexpressed YAP-TFE3-GFP.
**F)** Chart showing the photobleach-corrected survival distribution of Halo-TEAD1 molecules in nuclei of HeLa cells in the presence or absence of YAP-TFE3-GFP, with bi-exponential fitting (solid lines). Dotted lines are ± 99% CI.
**G and H)** Charts showing the mean long and short DNA dwell times (G) and relative proportion of long and short DNA dwell times (H) for TEAD1 molecules in the presence or absence of overexpressed YAP-TFE3-GFP in HeLa cell nuclei, as derived from the bi-exponential function. Dwell times are calculated as 1/k from the bi-exponential fit and displayed ± 95% CI.

**MOVIES**

**Movie S1.** Fast single molecule imaging of Halo-TEAD1 and Halo-YAP in MCF10A cells.

**Movie S2.** Slow single molecule imaging of Halo-TEAD1 and Halo-YAP in MCF10A cells.

**Movie S3**. Single molecule dynamics of Halo-tagged TEAD1, TEAD1$^{\Delta DBD}$, and TEAD1$^{Y421H}$ in HeLa cells.

**Movie S4.** Single molecule dynamics of Halo-TEAD1 in nuclear YAP-GFP condensates.
